# Defining redundancy in the stickers and spacers of the cell-cell junction protein Canoe’s intrinsically disordered region

**DOI:** 10.64898/2026.09.28.754968

**Authors:** Corbin C. Jensen, Renick Wiltshire, Sarah E. Clark, Jeremy Lamb, Kevin C. Slep, Mark Peifer

## Abstract

Cell-cell adherens junctions (AJs) and their dynamic cytoskeletal linkage power morphogenesis. AJs are enormous complexes with hundreds of proteins linked by multivalent interactions. Like other biomolecular condensates, intrinsically disordered regions (IDRs) in junctional proteins play important roles in AJ assembly and function, using spacer elements to span distances, and stickers to engage targets. To define molecular mechanisms, we need to define the functional units within IDRs. Drosophila Canoe, homolog of human Afadin, is our model. Canoe mediates morphogenesis and has an extensive IDR, with two conserved F-actin-binding stickers and two poorly conserved spacers. We combined biochemical, genetic and cell biological approaches to define the function of these IDR elements. While no single element is essential, deleting the full IDR essentially eliminates Canoe function. By scrambling the amino acid sequence of the spacers, we find that length and composition are more important than amino acid sequences, though sequences in the C-terminal spacer affect Canoe localization. Finally, we test redundancy of the F-actin-binding stickers. Deleting both reduces but does not eliminate viability, and sensitized assays reveal their redundant roles. These data reveal the robustness of IDRs.

## Introduction

While textbooks often portray simple protein complexes driving key cellular events, many are carried out by biomolecular condensates, massive complexes containing hundreds to thousands of proteins. Condensates concentrate molecules required for diverse events ranging from ribosome assembly to transcriptional regulation to cell-cell and cell-matrix adhesion (Holehouse and Kragelund, 2024; Rouaud et al., 2020; Sun et al., 2022). Multivalent protein interactions are key to their assembly and function. Some interactions are mediated by folded protein domains, but others involve intrinsically disordered regions (IDRs; (Holehouse and Kragelund, 2024)). One key challenge is to define the molecular mechanisms mediating the function of these multiprotein complexes.

IDRs do not fold on their own but can mediate low and high affinity interactions with folded protein domains or other IDRs. Broadly, IDRs include two different types of sequence: “Stickers” and “Spacers” (Ginell and Holehouse, 2023). Stickers or short linear motifs (SLiMs) serve as more specific binding sites for other proteins (Cermakova and Hodges, 2023). These can include short, structured segments or segments that fold upon interaction with a protein partner (Arai et al., 2024; Devi et al., 2025). Stickers are separated by “spacers” that often are low complexity sequences. They can provide flexible spacing between stickers, and can mediate lower affinity interactions via properties like electrostatic and aromatic interactions, driven by amino acid composition and charge (Koyama et al., 2024; Yang et al.).

We focus on the adherens junction (AJ) complex. AJs mediate cell-cell adhesion and link to the actomyosin cytoskeleton during morphogenesis and homeostasis, powering cell shape change and ensuring tissue integrity (Perez-Vale and Peifer, 2020; Yap et al., 2018). At its center is the cadherin-catenin complex. Classical cadherins mediate adhesion via their extracellular domains, and link to the cytoskeleton via their intracellular domains and associated catenin proteins. However, the picture is more complex. First, rather than forming the textbook “belt” around the apical end of the cell, cadherins assemble into large puncta in both mammalian and Drosophila cells (Harris and Peifer, 2004; Troyanovsky, 2023; Yap et al., 2015). Puncta vary in size but molecule counting in Drosophila embryos revealed that each punctum contains on average 1500 cadherins (McGill et al., 2009). Second, cadherin-catenin complexes are only the core of the AJ, with dozens of other proteins localizing there (Rouaud et al., 2020), linking the core complex to the actin cytoskeleton and reinforcing connections under tension. In fact, super-resolution microscopy is beginning to suggest that the AJ is not a simple, homogeneous entity, but instead may include multiple adjacent substructures mediating the cytoskeletal interface (Schmidt et al., 2023).

Most AJ proteins have multiple protein domains and one or more IDRs (Rouaud et al., 2020; Sun et al., 2022). To understand AJ function, we need to dissect the structures of the underlying machines to define the mechanisms by which their complex structures mediate function. Drosophila Canoe (Cno) and its mammalian ortholog Afadin provide superb entry points. Unlike cadherins and catenins (e.g. Cox et al., 1996; Larue et al., 1994), Cno and Afadin are not essential for adhesion itself (Ikeda et al., 1999; Sawyer et al., 2009). Instead, they mediate many events requiring linkage of AJs to the F-actin cytoskeleton. For Cno, this includes initial AJ positioning during cellularization, mesoderm apical construction, germband convergent extension, dorsal closure, and head involution, as well as postembryonic events shaping the pupal eye (Boettner et al., 2003; Choi et al., 2013; Matsuo et al., 1999; Sawyer et al., 2011; Sawyer et al., 2009). Cno strengthens AJ:cytoskeletal linkages. In Cno’s absence, linkage is weakened and often fails when junctions are under tension. Tension further modulates Cno localization (Zheng et al., 2026). Afadin similarly mediates diverse events in embryonic and postembryonic development (Ikeda et al., 1999; Tanaka-Okamoto et al., 2014; Yamamoto et al., 2013; Yang et al., 2013; Zhadanov et al., 1999).

Cno and Afadin share a complex protein structure, with five N-terminal folded domains and a ∼1000 amino acid C-terminal IDR (Fig. 1A; Gurley et al., 2023). Some protein domains have known functions: the N-terminal RA domains each bind the activated small GTPase Rap1, and the PDZ domain binds E-cadherin (Ecad) and transmembrane junctional Nectin family proteins. Two conserved motifs in the IDR bind F-actin, and, in the case of mammalian Afadin, other motifs bind alpha-catenin and other proteins. We initially tested a simple hypothesis: Rap1 binding activates Cno, which then links Ecad and actin, strengthening AJ:cytoskeletal linkage (Perez-Vale et al., 2021). However, while deleting the Rap1 binding RA1 domain nearly eliminated protein function, animals lacking the PDZ domain or the C-terminal F-actin binding segment (FAB) were viable and fertile. Sensitized assays revealed these mutants do not retain full function, but these data emphasized the robustness provided by multivalent interactions. In fact, three of the five folded domains (RA2, the Dilute domain and the PDZ) are dispensable for viability (McParland et al., 2024a; McParland et al., 2024b).

**Figure 1.**
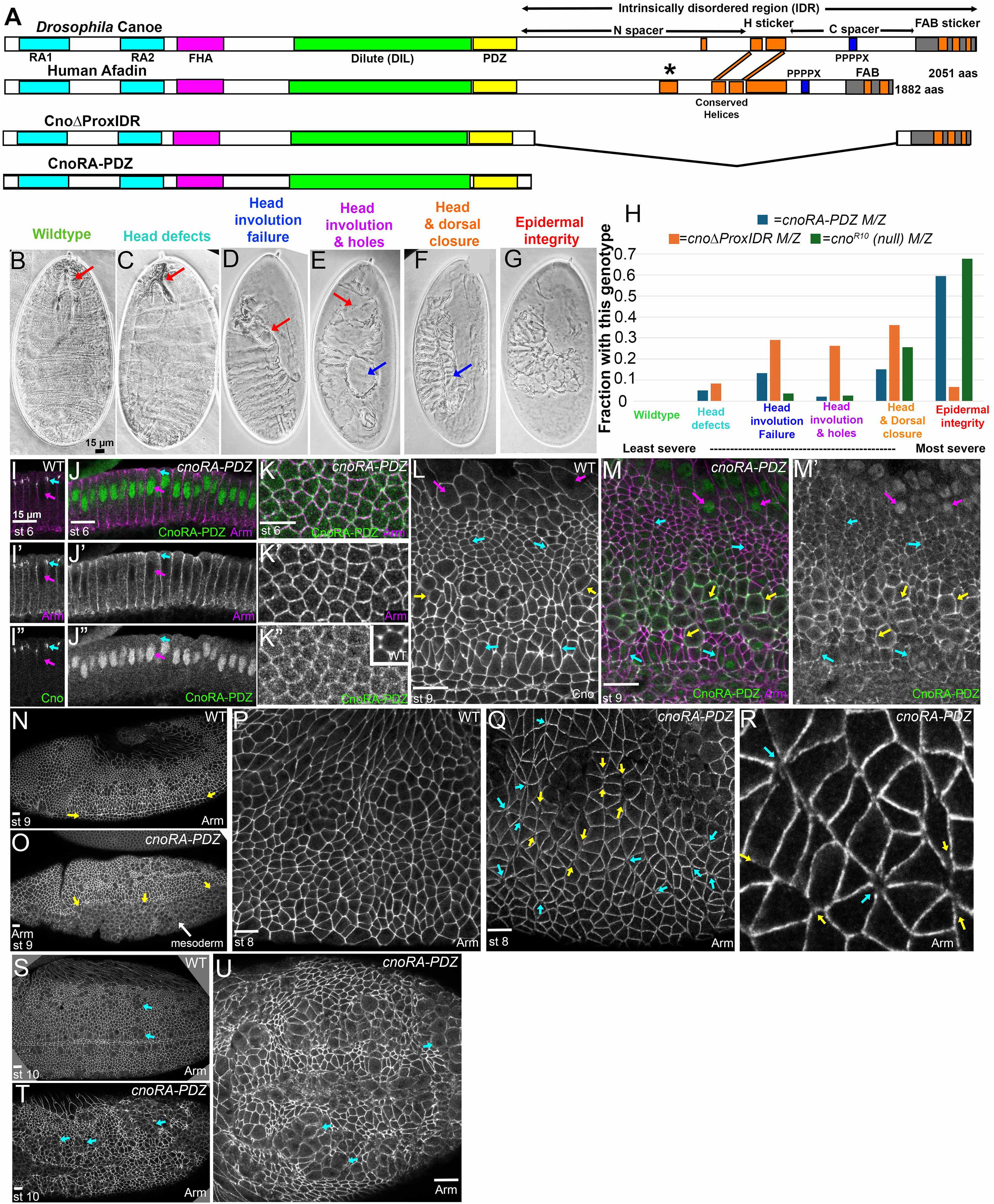
Removing Cno’s IDR disrupts localization and essentially eliminates protein function. A. Diagrams of Cno, Afadin and selected mutants. B-G. Cuticle phenotypic categories. Derived from Jensen et al. (2025). H. The maternal/zygotic cuticle phenotype of *cnoΔIDR* is substantially stronger than that of *cnoΔProxIDR* and is almost as strong as that of the *cno* null mutant. I-U. Embryos, genotypes, embryonic stages, and antigens indicated. I, J. Cross sections. I. Cno localizes to apical AJs (cyan arrow) and is absent from nuclei (magenta arrow). J. CnoΔIDR protein is almost absent from AJs (cyan arrow) and enriched in nuclei (magenta arrow). K. en face. CnoΔIDR protein localization to AJs is reduced and tricellular junction enrichment is lost. K inset, wildtype Cno, same stage. L. Wildtype Cno is enriched at AJs of non-mitotic cells (cyan arrows), reduced at the cortex of mitotic cells (yellow arrow), and absent from amnioserosal nuclei (magenta arrow). M. CnoΔIDR weakly localizes to AJs of non-mitotic cells (cyan arrows) but is elevated at the cortex of mitotic cells (yellow arrows) and enriched in amnioserosal nuclei (magenta arrows). N. Wildtype. Closed ventral furrow (arrows). O. *cnoΔIDR*. Wide open ventral furrow. P. Wildtype. AJs have few gaps. Q, R. *cnoΔIDR*. Examples of AJ gaps at rosette centers (yellow arrows) and aligned AP junctions (cyan arrows). R. Closeup. S. Wildtype. Mitotic cells round up and then rapidly return to columnar shape. T, U. Mitotic cells are slow to return to columnar shape (arrows). Arm accumulation is reduced.

Given this, we turned to Cno’s IDR. Combining sequence analysis, AlphaFold modeling, biochemical experiments and the literature allowed us to roughly divide the Cno/Afadin IDR into four segments, with two conserved stickers and two spacers (Fig 1A; (Jensen et al., 2025). The two conserved stickers in the IDR each can bind F-actin (Gong et al., 2025; Jensen et al., 2025): these include the C-terminal F-actin binding segment (FAB), and a central segment of two helices (H) which cryo-EM and biochemical analyses reveal stabilizes interactions between alpha-catenin and F-actin (Gong et al., 2025). These are separated from the PDZ domain and one another by two poorly conserved spacers (N and C). In mammalian Afadin, the N spacer contains a binding site for alpha-catenin’s M domain but this is not conserved in Drosophila (Gurley et al., 2023; Pokutta et al., 2008). The C spacer appears to carry a nuclear export signal, as deleting it leads to accumulation of both Cno and Afadin in nuclei (Jensen et al., 2025; Kuno et al., 2025). Deleting the entire proximal IDR reduced Cno localization to AJs and led to embryonic lethality, with strong disruption of morphogenesis, though detailed analysis suggested residual protein function (Jensen et al., 2025). A similar deletion of Afadin’s proximal IDR also led to reduced AJ localization and function (Kuno et al., 2025). However, to our surprise, smaller deletions revealed that no individual segment of the IDR is essential Galapon, including the conserved stickers (Jensen et al., 2025).

These data raised several important questions about the mechanisms by which complex IDRs mediate AJ protein function. We set out to answer those. First, we examined the significant residual function retained by CnoΔProxIDR (Jensen et al., 2025), by creating a mutant deleting the full IDR to see if retaining the FAB segment underlies the retained function in morphogenesis. Second, while we could individually delete the N and C spacers, deleting both strongly reduced protein function (Jensen et al., 2025). This raised the question of whether the amino acid sequence of two spacers is important, or if they simply provide “spacer” function. We tested this by scrambling the sequence of the two spacers, thus retaining their respective lengths and amino acid composition. Finally, we further explored the role of the F-actin binding stickers. We tested the hypothesis that they are individually dispensable because they are redundant in function, by deleting both stickers. Together, our experiments provide detailed mechanistic insights into how IDR’s mediate function in a junctional protein, revealing how the stickers and spacers combine to provide full function.

## Results

### Deleting the full IDR strongly reduces or eliminates Cno function

While our prior work revealed an important role for the proximal IDR in Cno function, we were surprised that no individual IDR segment (N, H, C, or FAB) is essential for viability (Jensen et al., 2025). Further, neither mutant in which multiple IDR segments were deleted completely eliminated function, as the embryonic phenotypes of *cnoΔProxIDR* and *cnoIDRHelixOnly* were weaker than those of the null mutant (Jensen et al., 2025). This raised the question of whether the entire IDR is essential for function. To test this, we generated a new mutant, deleting the entire IDR including the FAB, which we refer to as *cnoΔIDR* (Fig 1A, Fig S1). We retained the segment just C-terminal to the PDZ domain, as it is strongly conserved in flies and mammals (Gurley et al., 2023). Like our other mutants, CnoΔIDR is C-terminally GFP-tagged. This and the other new mutants below were confirmed by PCR and sequencing. We assessed protein accumulation by immunoblotting using antibodies to GFP (Fig S2). CnoΔIDR protein accumulated at levels somewhat higher than wildtype CnoGFP.

As expected, *cnoΔIDR* was lethal over a *cno* null allele (*cno^R2^;* n=635). Thus, to analyze its function in embryonic development, we generated maternal-zygotic mutants using the FLP/FRT/*ovo^D^* approach (Chou et al., 1993), crossing females with germlines homozygous for *cnoΔIDR* with *cnoΔIDR/+* males. 55% of the embryos died before hatching, consistent with full lethality of maternal-zygotic mutants and penetrant zygotic rescue. We then assessed CnoΔIDR’s function in morphogenesis by analyzing the cuticles of maternal-zygotic mutants (Fig. 1B-G, arranged least severe to most severe). This provides our most sensitive assay, allowing us to assess completion of multiple morphogenetic events requiring Cno, including head involution (Fig 1B vs C, D), dorsal closure (Fig 1B vs E, F), and ventral epidermal integrity (Fig. 1G). Epidermal integrity is strongly disrupted in *cno* maternal-zygotic null mutants while the epidermis in *cnoΔProxIDR* is much less affected (Fig. 1H; Gurley et al., 2023; Jensen et al., 2025; Sawyer et al., 2009). *cnoΔIDR* maternal-zygotic mutants had penetrant defects in head involution and dorsal closure and the majority also had defects in ventral epidermal integrity (Fig. 1H; 59%). This was substantially stronger than the phenotype of *cnoΔProxIDR* (7% epidermal integrity defects (Jensen et al., 2025) and nearly as severe as the defects seen in maternal-zygotic null mutants (68%; (Gurley et al., 2023). This suggests removing the IDR essentially eliminates Cno function and reveals that the FAB segment is responsible for the retained function of CnoΔProxIDR.

### CnoΔIDR protein has reduced junctional localization and accumulates in nuclei

Wildtype Cno localizes to apical AJs (Fig, 1I, K” inset). This localization is quite robust, as deleting the RA domains, the Dilute domain, the PDZ domain or the C-terminal FAB do not disrupt it (McParland et al., 2024a; McParland et al., 2024b; Perez-Vale et al., 2021). The only mutants we examined with reduced localization to AJs either deleted the entire proximal IDR or removed both its N and C-terminal spacers (Jensen et al., 2025). In addition, these two mutant proteins also re-localized to nuclei. To explore the role of the full IDR, we examined CnoΔIDR localization.

We first looked at the onset of gastrulation. At this stage cadherin-catenin complexes, visualized with Arm antibody, localize all along the lateral interface and are enriched at apical AJs, while Cno is restricted to AJs (Fig. 1I, cyan arrow). In contrast, AJ localization of CnoΔIDR was strongly reduced and the protein instead localized to nuclei (Fig. 1J, cyan vs magenta arrows). When viewed en face, weak junctional localization of CnoΔIDR was seen (Fig 1K, K”), but the obvious tricellular junction (TCJ) enrichment of wildtype Cno (Fig 1K”, inset; (Bonello et al., 2018) was not apparent. This pattern of weak junctional localization in non-mitotic cells continued throughout embryonic morphogenesis. For example, at stage 9 in wildtype, Cno is strongly enriched at AJs of non-mitotic cells (Fig. 1L, cyan arrows), while AJ accumulation is somewhat reduced as cells round up for mitosis (Fig. 1L, yellow arrows). In contrast, CnoΔIDR localization was relatively weak in non-mitotic cells (Fig. 1M, cyan arrows), but, like CnoΔProxIDR (Jensen et al., 2025), CnoΔIDR was cortically enriched in cells rounding up for mitosis (Fig. 1M, yellow arrows). Nuclear localization remained prominent, as is seen in amnioserosal cells (Fig. 1L vs M, magenta arrows). Thus, CnoΔIDR retains some ability to localize to AJs, but, like CnoΔProxIDR, this localization is reduced and replaced by strong nuclear accumulation, due, we suspect, to deletion of a nuclear export signal in segment C.

### Deleting the full IDR strongly impairs Cno function during morphogenesis

The first morphogenetic movement requiring Cno is apical constriction of mesodermal cells, and *cno* null mutants have penetrant defects in ventral furrow closure (Sawyer et al., 2009). *cnoΔIDR* maternal-zygotic mutants, identified by the lack of staining with our C-terminal Cno antibody, shared this defect.16/17 stage 7-9 embryos had an open ventral furrow. The phenotypes were quite severe, with 10/17 (59%) exhibiting a wide-open ventral furrow (Fig. 1N vs O) while 6/17 had more moderate defects with the anterior end open (Table 1). This is substantially more severe than what we observed in *cnoΔProxIDR* mutants where only 22% had a wide-open ventral furrow while in 19% the furrow was fully closed (Jensen et al., 2025).

**Table 1.** Defects in Ventral furrow invagination.

| Genotype | VF OK | Mild defects | Moderate defects | Wide open | Total | Percent defective |
| --- | --- | --- | --- | --- | --- | --- |
| +/*cno*<sup>R2</sup> @ | 48 | 3 | 0 | 0 | 51 | 6% |
|  | 94% | 6% | 0 | 0 |  |  |
| *cnoIDRΔN*/*cno*<sup>R2</sup> # | 29 | 5 | 1 | 1 | 36 | 19% |
|  | 80% | 14% | 3% | 3% |  |  |
| *cnoIDRΔC*/*cno*<sup>R2</sup> # | 21 | 13 | 5 | 5 | 44 | 52% |
|  | 48% | 30% | 11% | 11% |  |  |
| *cnoIDRScramble*/*cno*<sup>R2</sup> | 22 | 11 | 4 | 0 | 37 | 40% |
|  | 59% | 30% | 11% | 0% |  |  |
| *cnoΔProxIDR* maternal/zygotic # | 6 | 10 | 8 | 7 | 31 | 81% |
|  | 19% | 32% | 26% | 23% |  |  |
| *cnoIDRHelixOnly* maternal/zygotic # | 18 | 23 | 33 | 8 | 82 | 78% |
|  | 22% | 28% | 40% | 10% |  |  |
| *cnoIDRΔHFAB* maternal x heterozygous males | 30 | 5 | 5 | 7 | 47 |  |
|  | 64% | 11% | 11% | 15% |  |  |
| *cnoIDRΔHFAB* adjusted for paternal rescue | 7 | 5 | 5 | 7 | 24 |  |
|  | 29% | 21% | 21% | 29% |  |  |
| *cnoIDRΔH*/*cno*<sup>R2</sup> # | 52 | 7 | 0 | 0 | 59 | 12% |
|  | 0.88135593 | 0.11864407 | 0 | 0 |  |  |

@ Data is from McParland et al. 2024 # Data is from Jensen et al., 2025

As the germband elongates, cells intercalate using planar polarized localization of junctional and cytoskeletal proteins. Cno stabilizes AJs under tension and, in its absence, AJs are destabilized, particularly at aligned anterior-posterior (AP) borders and the centers of rosettes (Sawyer et al., 2011). While wildtype AJs are continuous (Fig. 1P), *cnoΔIDR* maternal-zygotic mutants exhibited strong defects in AJ integrity at aligned AP borders (Fig. 1Q,R, yellow arrows) and rosette centers (Fig. 1Q,R, cyan arrows). These gaps were large, like those seen in *cno* null mutants or after strong RNAi (Manning et al., 2019; Sawyer et al., 2011), and thus unlike the smaller gaps we observed in many of our other mutants, including *cnoΔProxIDR* (Jensen et al., 2025). In wildtype, cells round up during division (Fig. 1S, arrows) but then rapidly resume columnar architecture, while *cno* null mutant cells have difficulty resuming columnar cell shape after mitosis. This is particularly problematic in the ventral epidermis (Sawyer et al., 2009). Like these strong mutants, *cnoΔIDR* maternal-zygotic mutants also had persistent rounded up cells in the ventral epidermis, with weakened junctional Arm (Fig. 1T,U, arrows). We suspect persistence of this leads to the ventral defects seen in the cuticles above. Thus, deleting the full IDR essentially eliminates Cno function, with *cnoΔIDR* maternal-zygotic mutants having phenotypes stronger than *cnoΔProxIDR,* and close to or the same as *cno* null mutants, and the FAB is responsible for the retained function of *cnoΔProxIDR*.

### Scrambling the IDR spacer sequences does not lead to lethality but impairs full wildtype function

Our exploration of Cno’s IDR revealed surprising redundancy, with no single segment, when deleted, substantially reducing Cno function or leading to embryonic lethality (Jensen et al., 2025). Two segments, termed H and the FAB, are “stickers”, carrying experimentally-verified F-actin binding sites conserved between Cno and Afadin (Fig 2A; (Jensen et al., 2025). The other two segments, termed N and C, appear to have “spacer” function, as they are low complexity sequences that are poorly conserved between flies and mammals (Fig. 2A; Gurley et al., 2023). Mutants individually lacking the N or C spacers (Fig 2A) are viable and fertile, though sensitized assays revealed they are not fully wildtype in function. However, simultaneously deleting N and C (CnoIDRHelixOnly; Fig 2A) impairs Cno localization to AJs and substantially reduces Cno function, leading to embryonic lethality and major morphogenesis defects (Jensen et al., 2025).

**Figure 2.**
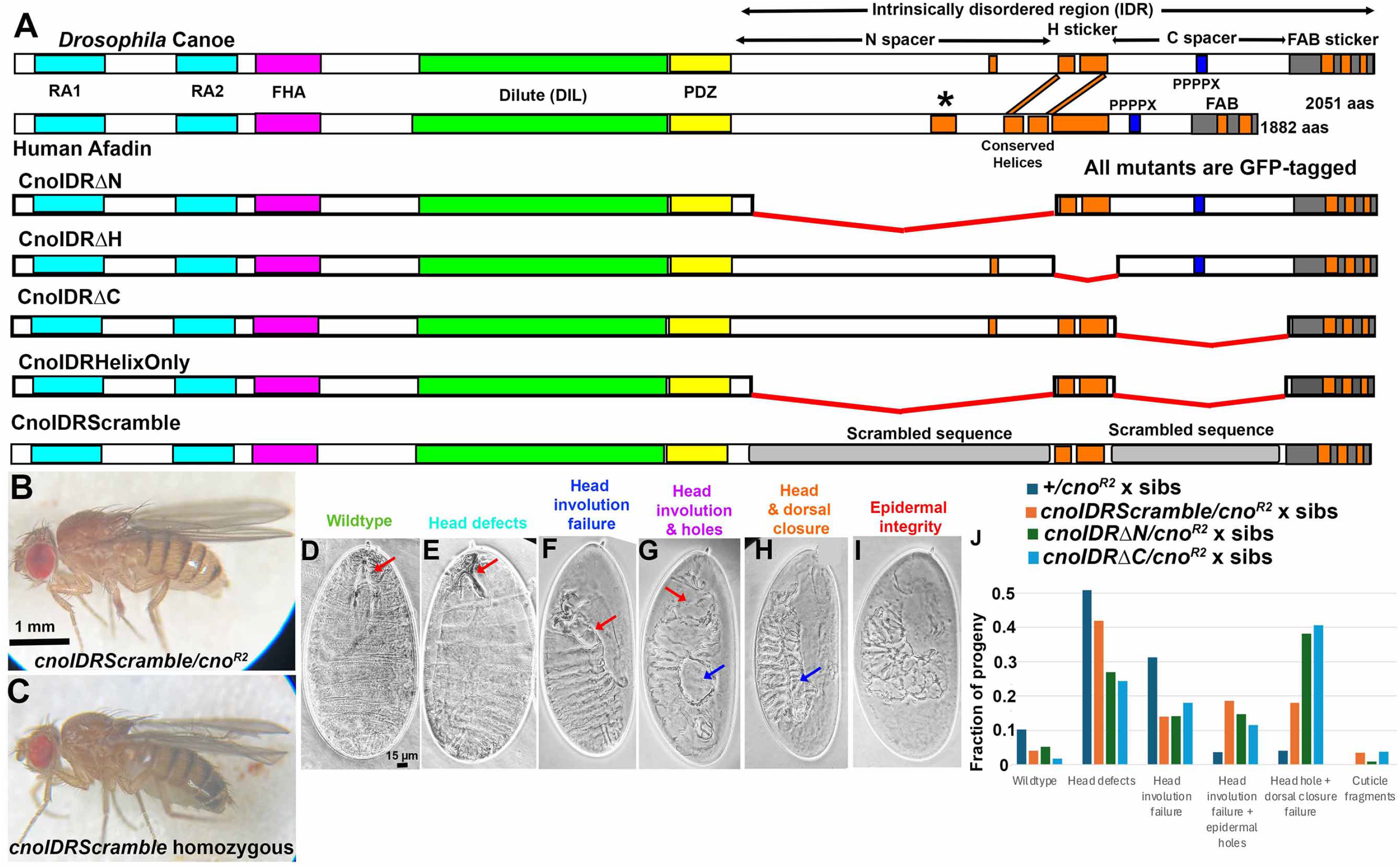
Scrambling the sequence of the IDR spacers only subtly reduced Cno function. A. Diagrams of selected mutant. B. *cnoIDRScramble/cnoR2* adults are viable. C. *cnoIDRScramble* homozygous adults are viable and fertile. D-I. Cuticle phenotypic categories. J. Quantification of cuticle phenotypes. CnoIDRScramble protein does not provide fully wildtype function, but in this assay is somewhat more functional than CnoIDRΔN and CnoIDRΔC.

We thus asked whether N and C carry functional sequence motifs, or whether their importance is solely in their length and sequence composition, by generating *cnoIDRScramble* (Fig. 2A). In this mutant, we left the F-actin binding stickers intact but scrambled the amino acid sequences of spacer segments N and C using the Shuffle Protein tool in the Sequence Manipulation Suite (Stothard, 2000). This randomly shuffled each protein sequence while retaining the same amino acid composition and length—the scrambled sequences continue to be predicted as disordered (Fig S1). We then replaced *cno* at its endogenous locus with this mutant, which was C-terminally GFP-tagged. Immunoblotting revealed a GFP-tagged protein of the appropriate size (Fig S2), accumulating at levels slightly higher than wildtype.

We first asked whether *cnoIDRScramble* mutants were viable. To do so and to avoid effects of other mutations on the chromosome, we crossed them to animals carrying a protein null allele, *cno^R2^*. *cnoIDRScramble/cno^R2^*flies were viable to adulthood and produced viable adult progeny (Fig. 2B). However, adult viability was reduced by two-thirds (10.8% rather than the 33% expected from wildtype (n=213; Balancer chromosome homozygotes are lethal). After outcrossing, we obtained homozygous mutants (Fig. 2C). This rules out an essential role in viability for any sequence motifs in the N or C spacers but the reduced viability suggested CnoIDRScramble protein does not provide full wildtype function.

Most *cno* mutants deleting individual folded protein domains or IDR segments were also viable as heterozygotes over *cno^R2^*, and we thus developed a sensitized assay to determine whether they retained fully wildtype function (Perez-Vale et al., 2021). To do so, we cross males and females heterozygous for our new allele and the null allele *cno^R2^.* All progeny lack wildtype Cno and have reduced levels of maternally contributed mutant protein, with 25% of embryos homozygous for our new mutant, 50% heterozygous for our new mutant and *cno^R2^,* and 25% homozygous for *cno^R2^. cno^R2^* zygotic mutants are embryonic lethal, and thus in the baseline cross of *+/cno^R2^* males and females, 25% of embryos die. We then used cuticle preps to assess completion of three events requiring Cno: dorsal closure, head involution, and epidermal integrity (Sawyer et al., 2009). Maternally contributed *cno* mRNA and protein is sufficient for early morphogenetic movements, so *cno^R2^/cno^R2^*mutants only have defects in the final morphogenetic movement, head involution (as in Fig. 2D vs E,F; quantified in 2J; (Gurley et al., 2023).

We used this assay to examine progeny of *cnoIDRScramble/cno^R2^*flies. Lethality was slightly elevated (30.3% versus the expected 25%), similar to *cnoIDRΔN* (34% lethality), and considerably less severe than *cnoIDRΔC* (53% lethality;(Jensen et al., 2025). *cnoIDRScramble/cno^R2^* progeny had more severe cuticle phenotypes than the +/*cno^R2^*standard, with 21% of the embryos having complete failure of dorsal closure and head involution, compared to ∼4% of the progeny of *+/cno^R2^* males and females (Fig. 2H, I, J). This was less severe than either *cnoIDRΔN* or *cnoIDRΔC,* where 38% or 41% of embryos have both dorsal closure and head involution failure, respectively (Jensen et al., 2025). Thus, while CnoIDRScramble protein rescues viability, it does not provide fully wildtype function. However, it is much more functional than a mutant deleting both spacers, as CnoIDRHelixOnly, is embryonic lethal with strong morphogenesis defects. Thus, restoring the “spacer” function of segments N and C is sufficient to restore viability and much of Cno’s function, with the modest reduction in function likely due to a requirement for specific sequences in the spacers.

### CnoIDRScramble protein localizes to both AJs and nuclei, and tension-sensitive AJ recruitment is diminished

The IDR, as a whole, plays a role in Cno localization to apical AJs and enrichment at AJs with elevated tension, including TCJs, as we observed above for CnoΔIDR and previously for CnoΔProxIDR and CnoIDRHelixOnly (Jensen et al., 2025). AJ enrichment of these mutants is reduced, TCJ enrichment is lost, and they instead accumulate in nuclei. However, no individual IDR segment (N, H, C, or FAB) is essential for AJ localization (Jensen et al., 2025; Perez-Vale et al., 2021), and neither H nor the FAB play a role in TCJ enrichment. However, while CnoIDRΔN and CnoIDRΔC accumulate at normal levels in AJs, deleting the N segment reduces TCJ enrichment and deleting the C segment nearly eliminates it. Surprisingly, deleting segment C also led to nuclear accumulation alongside AJ localization (Jensen et al., 2025).

We thus tested whether these roles of the N and C segments were conferred by length and sequence composition, or whether there were specific important sequences in these segments. CnoIDRScramble localized to AJs from cellularization through the end of morphogenesis, like wildtype (e.g. Fig 3A, B). While cadherin-catenin complexes localize all along the lateral membrane, CnoIDRScramble, like wildtype Cno, was enriched in apical AJs (Fig. 3C, D, yellow vs cyan arrows). However, CnoIDRScramble also localized to nuclei (Fig 3C vs D, magenta arrows, Fig 3B’). Next, we examined TCJ enrichment. Wildtype Cno is strongly enriched TCJs in response to tension, relative to bicellular junctions (Fig. 3E, magenta vs cyan arrows; (Yu and Zallen, 2020) (Perez-Vale et al., 2021). TCJ enrichment of CnoIDRScramble was strongly reduced but not eliminated (Fig. 3F, G magenta vs cyan arrows;1.54 fold-enriched relative to 2.94 fold in wildtype). In both properties, CnoIDRScramble resembled CnoIDRΔC, suggesting that the precise sequence of segment C is important for both nuclear exclusion and TCJ enrichment.

**Figure 3.**
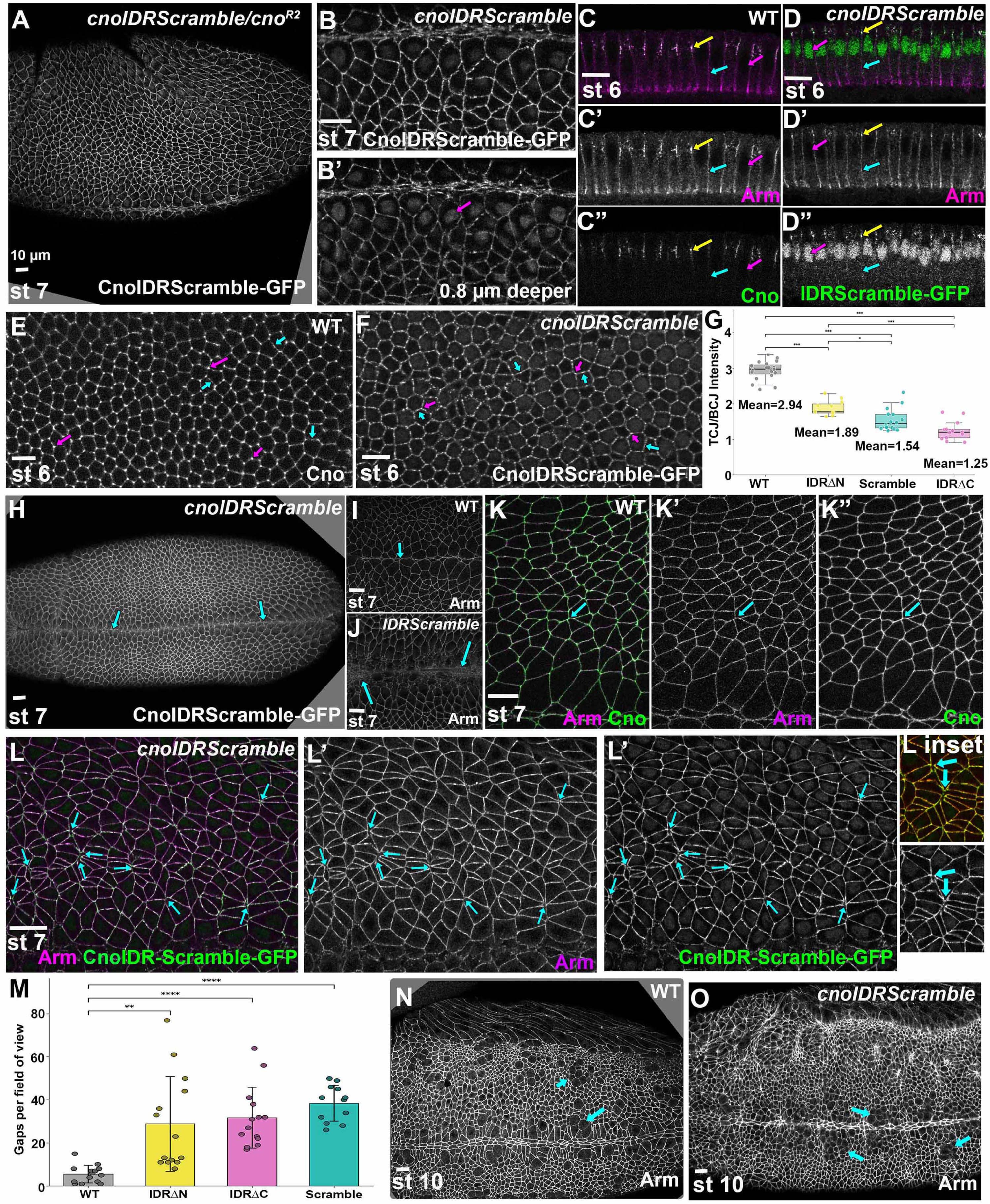
CnoIDRScramble localizes to both AJs and nuclei and has mild deficits in function. A-F, H-L, M, N. Embryos, genotypes, embryonic stages, and antigens indicated. A, B. *en face*. C, D. cross sections. C. Wildtype Cno is restricted to apical AJs, (yellow arrow) and absent from lateral borders (cyan arrow) and nuclei (magenta arrow). D. CnoIDRScramble is also enriched at apical AJs (yellow arrow) relative to lateral borders (cyan arrow) but accumulates in nuclei (magenta arrow). E, F. While wildtype Cno is enriched at TCJs relative to bicellular borders (E, magenta vs cyan arrows), TCJ enrichment of CnoIDRScramble is reduced. F. TCJ enrichment. H-J. Most *cnoIDRScramble* mutants complete ventral furrow invagination (H, arrow) like wildtype (I), and if defects are seen they are mild (J, arrows). K. Wildtype. AJs have very few gaps. L. *cnoIDRScramble* has a somewhat elevated number of small gaps. (cyan arrows). M. Gap frequency. N. Wildtype cells round up for mitosis (arrows) and then resume columnar shapes. O. Some *cnoIDRScramble* mutants have delays in cells resuming columnar shapes (arrows).

### *cnoIDRScramble* mutants have reduced integrity of AJs under tension

We next explored how scrambling the spacers affected embryonic morphogenesis. We first examined ventral furrow invagination, using as a baseline progeny of +/*cno^R2^*parents, 6% of which have mild defects in this process (McParland et al., 2024a); Table 1). Most progeny of *cnoIDRScramble/cno^R2^* flies completed ventral furrow invagination (Fig. 3H; 59%; 22/37 embryos), and if defects were seen they were always mild (Fig 3J; 30%; 11/37) or moderate (11%; 4/37). These defects were more severe than those of *cnoIDRΔN/cno^R2^* progeny (81% completed invagination) but less severe than defects seen in *cnoIDRΔC/cno^R2^*progeny (48% completed furrow formation and 11% had a wide-open ventral furrow; (Jensen et al., 2025).

We next examined phenotypes during germband extension, when Cno is enriched at and stabilizes AJs under elevated tension (Sawyer et al., 2011; Yu and Zallen, 2020). Wildtype AJs have very few gaps and those that are present are small (Fig. 3K). Among *cnoIDRScramble/cno^R2^* progeny gap frequency was elevated along shrinking AP borders and at rosette centers (Fig.3L, arrows), with an average of 36 gaps per field of view (n=14 stage 7 and 8 embryos; Fig 3M) vs the 15 gaps per field we observed in our +/*cno^R2^*control (Jensen et al., 2025). *cnoIDRScramble/cno^R2^* gap frequency was roughly similar to that in *cnoIDRΔC/cno^R2^* progeny (32 gaps/field), slightly higher than was seen in *cnoIDRΔN/cno^R2^* progeny (29 gaps/field), but much lower than what we saw in *cnoIDRHelixOnly* maternal/zygotic mutants (77 gaps/field; (Jensen et al., 2025). In *cno* null or *cnoΔIDR* maternal-zygotic the ventral epidermis is strongly destabilized (e.g. Fig 1U), leading to holes in the cuticle. We did not observe this in *cnoIDRScramble/cno^R2^* progeny. However, in milder mutants we previously observed delays in the return of dividing cells to columnar shape. Similarly, a subset of *cnoIDRScramble/cno^R2^* progeny exhibited mild delays in this (Fig. 3O, arrows; 5/13 stage 9-10 embryos), in contrast to cells in wildtype that round up to divide (Fig 3N, arrows) and rapidly resume columnar architecture. Thus, CnoIDRScramble protein retains substantial but not full wild-type function, and the nature and level of its defects most closely resemble those of CnoIDRΔC. This suggests spacer length rather than sequence is most important for Cno localization and function, but that sequences in both spacers influence enrichment at TCJs and sequences in segment C confer nuclear exclusion.

### *cnoIDRScramble* mutants have modest defects in pupal eye development

Cno also regulates postembryonic development. The developing eye provides a place to explore the function of *cno* mutants in complex cell shape changes. Each ommatidium has four cone cells, two primary (1°) cells, six secondary (2°), three tertiary (3°) cells, and three bristles, each with a characteristic position, shape and arrangement (Fig 4A; Johnson, 2021). Clones of cells null for *cno* severely disrupt this epithelium (Walther et al., 2018), while milder mutants, including those we examined in our structure-function studies (e.g. Jensen et al., 2025), have more subtle defects, thus providing an alternate place to assess phenotypic severity. We examined *cnoIDRScramble/cno^R2^*eye discs at 40 hours after puparium formation, visualizing Ecad and N-cadherin, and using the GFP-channel to visualize mutant Cno. We used an established scoring system (Johnson and Cagan, 2009) to score defects in stereotyped cell arrangement, including defects in the number and arrangement of cone, 1°, and 3° cells, and bristles per ommatidium (Table S2). For each genotype we scored 10-14 eye discs with 88-137 total ommatidia. Wildtype eyes, those of wildtype GFP-tagged Cno, or of *cno^R2^* heterozygotes have occasional defects: ∼0.5 defects per ommatidium (Fig. 4B, F, Table S1; Perez-Vale et al., 2021). In contrast, *cnoIDRScramble/cno^R2^* had more frequent defects, with 2.50 defects per ommatidia (Fig. 4C, F, Table S1). This tended slightly less severe than observed in *cnoIDRΔN/cno^R2^* (2.96 defects per ommatidium; Fig. 4D, F, Table S1), but didn’t reach statistical significance. It was substantially less severe than observed in *cnoIDRΔC/cno^R2^* (3.80 defects per ommatidium; Fig. 4C, F, Table S1; (Jensen et al., 2025). Since *cnoIDRScramble/cno^R2^* were viable to a stage where we could score eyes, unlike CnoIDRHelixOnly, the spacer function is most relevant but clearly sequence also plays a role.

**Figure 4.**
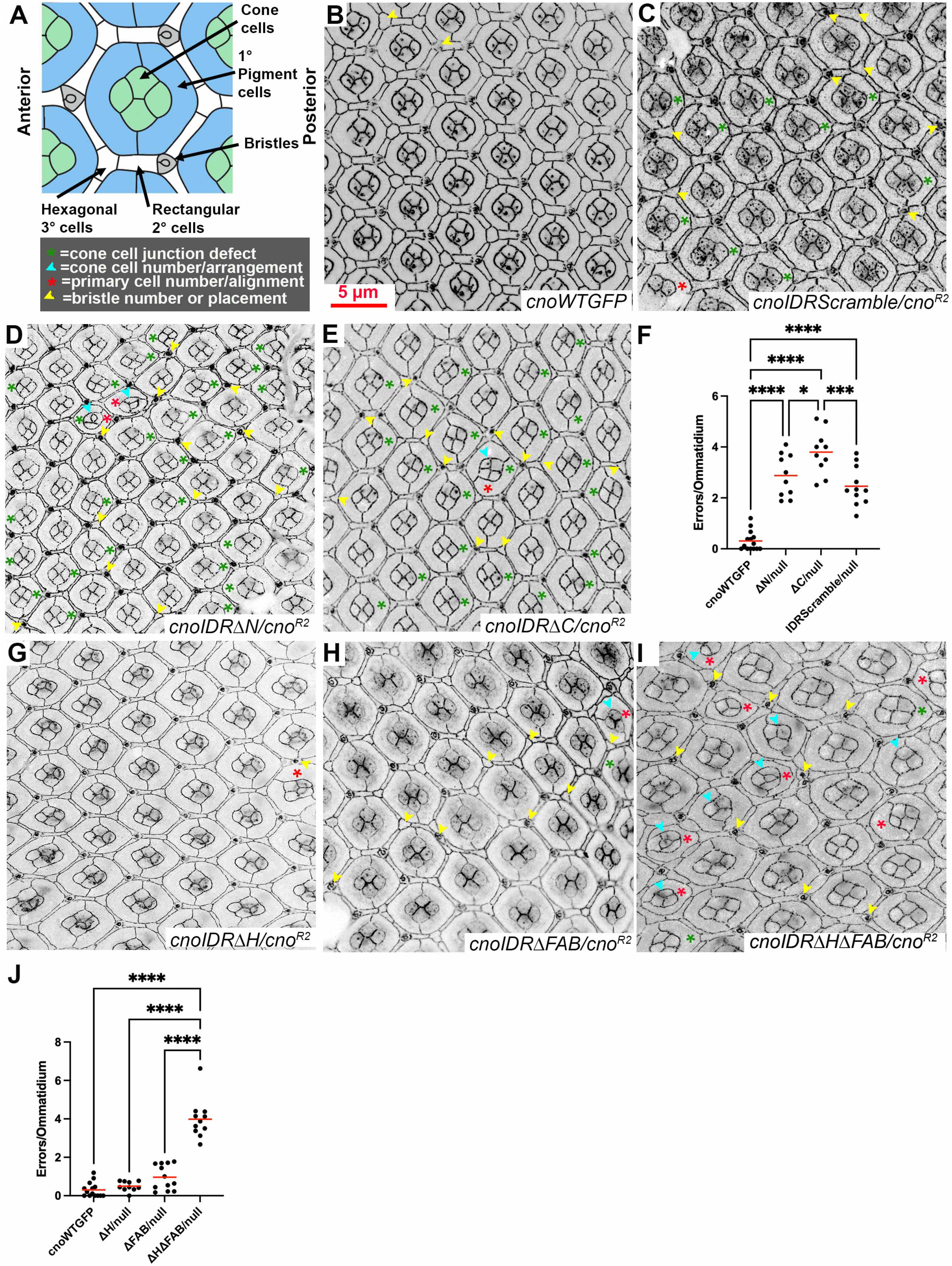
Assessing pupal eye morphogenesis in *cnoIDRScramble* and *cnoIDRΔHΔFAB* mutants. A. Diagram of pupal ommatidium and scoring criteria. B-E, G-I. Representative pupal eyes of the indicated genotypes, imaged to visualize Ecad, N-cadherin and GFP-tagged Cno variants. F, J. Quantitative comparison of defects per ommatidium. *cnoIDRScramble/cno^R2^*. C. *cnoIDRScramble/cno^R2^* mutants have a moderate level of defects. I, J. *cnoIDRΔHΔFAN/cno^R2^* defect frequency is much higher than that of either single deletion.

**Figure 5.**
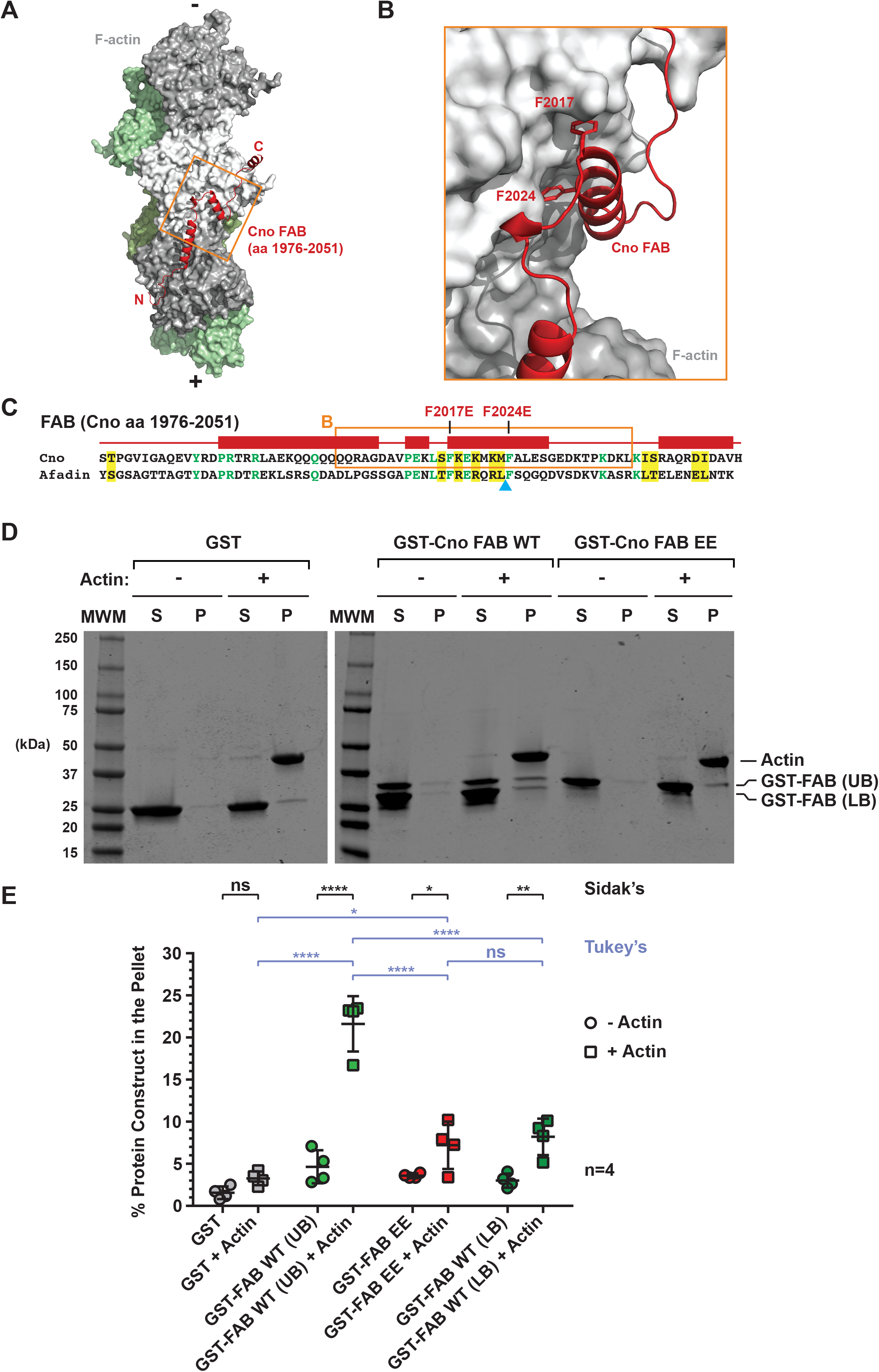
Cno’s FAB uses conserved hydrophobic residues to engage F-actin. A. AlphaFold model of Cno’s FAB (residues 1976-2051, red) bound to F-actin (one protofilament in grey tones, the other in green tones; Jensen et al. (2025)). B. Zoom of the region boxed in orange in A and C, after a 30° rotation about the y-axis and a 30° rotation about the x-axis relative to A. F2017 and F2024 are shown in stick format. C. Sequence alignment of Cno and rat Afadin FABs with cross-species identity in green and similarity in yellow. AlphaFold predicted helices=red rectangles above the alignment. Cyan arrowhead below the alignment: the Cno FAB C-terminal truncation product mapped in Jensen et al. (2025). D) Sedimentation analysis of GST, GST-Cno FAB (aa 1976-2051) WT and the F2017E/F2024E double mutant (EE) in the absence or presence of F-actin, n=4. E. Quantification. Statistical significance calculated using two-way ANOVA followed by either Sidak’s or Tukey’s multiple comparisons test.

### Tests support the AlphaFold model for how the FAB binds F-actin

We next turned to the IDR’s F-actin binding activity. Two regions can bind F-actin: the helical (H) region between IDR-N and IDR-C, and the C-terminal, conserved F-actin binding (FAB) region. Biochemical and structural biology analysis of the H region in mammals revealed it stabilizes quaternary interactions between α-catenin’s actin-binding domain and F-actin (Gong et al., 2025), and F-actin binding assays and AlphaFold modeling suggests the Drosophila H region plays a similar function (Jensen et al., 2025). The Drosophila FAB (residues 1967-2051) binds F-actin, and AlphaFold modeling suggested a potential mode of interaction involving a helix-turn-helix motif binding to actin subdomain I, using a conserved set of charged and hydrophobic residues (Fig 6A-C, described in (Jensen et al., 2025). In our published F-actin co-sedimentation assay using purified FAB protein, we noted the presence of a C-terminal degradation product with weaker co-sedimentation activity. Mass spectroscopy revealed that it lacked part of the helix-turn helix motif (Fig 6C, see cyan arrowhead), suggesting this contributes to F-actin binding. To further test this model, we generated an N-terminal GST-fusion FAB construct (GST-FAB WT) focused on the most highly conserved FAB region, residues 1976-2051. We also generated a double point mutant, targeting two conserved phenylalanine residues, F2017 and F2024, that the AlphaFold model predicted interact directly with F-actin (Fig 6B,C). Both phenylalanines were mutated to glutamate to introduce repulsive charges. We then performed F-actin co-sedimentation assays with these constructs and a GST control, mixing proteins with polymerized F-actin, pelleting F-actin filaments, and assessing how much is found in the pellet with F-actin. GST alone exhibits only background levels of co-sedimentation. GST-FAB WT showed significant co-sedimentation activity, on par with the larger FAB construct (residues 1967-2051) tested in Jensen et al., (2025). A smaller C-terminal degradation product of the GST-FAB WT construct was once again present and had weaker binding activity relative to the full-length construct (Fig 6D,E; compare GST-FAB WT upper band (UB) and lower band (LB)), highlighting the importance of determinants in the FAB’s C-terminal region. In contrast, the GST-FAB F2017E/F2024E (GST-FAB EE) mutant construct showed relatively weak F-actin co-sedimentation activity, on par with the GST-FAB WT truncated protein (Fig 6D,E), suggesting these conserved hydrophobic residues are involved in F-actin binding, as predicted by the AlphaFold model.

**Fig 6.**
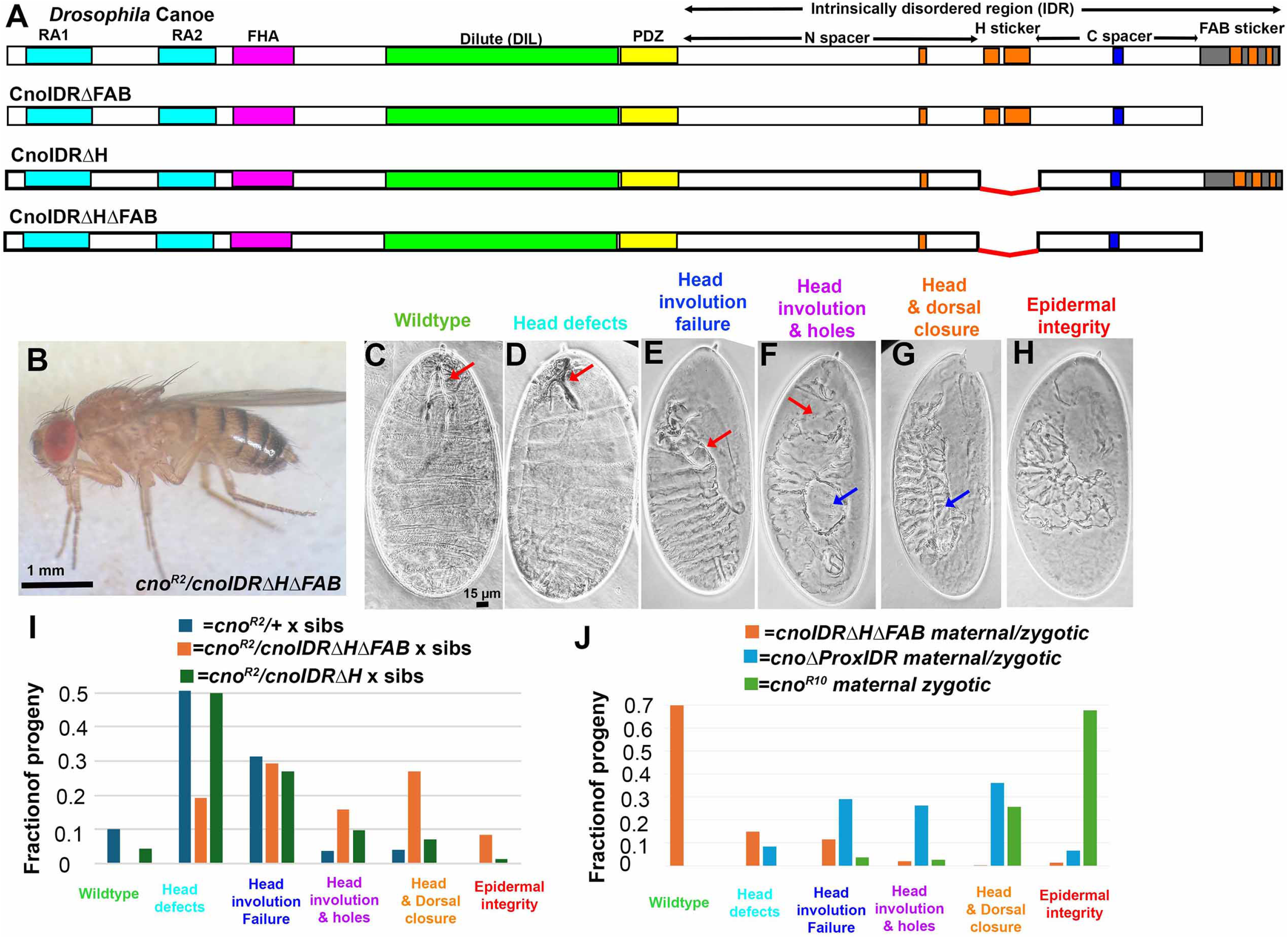
Deleting both F-actin binding stickers strongly reduces but does not eliminate adult viability and leads to relatively modest defects in embryonic morphogenesis. A. Diagrams of selected mutants. B. Zygotic *cnoIDRΔHΔFAB/cno^R2^* adult. C-H. Cuticle phenotypic categories. I. Sensitized assay reveals CnoIDRΔHΔFAB does not reveal fully wildtype function. J. Analysis of maternal/zygotic *cnoIDRΔHΔFAB/cno^R2^* mutants reveals modest defects in morphogenesis.

### Deleting both F-actin binding stickers strongly reduces but does not eliminate adult viability

Both the H region and the FAB are individually dispensable for viability, though they are important for full wildtype function (Jensen et al., 2025). We hypothesized that the two F-actin binding sites might be redundant in function, with each compensating for the loss of the other. To test this, we generated a mutant, *cnoIDRΔHΔFAB*, deleting both stickers but leaving the N and C spacers intact (Fig 6A; Fig S1). Once again, we assessed accumulation of an appropriately sized GFP-tagged protein by immunoblotting (Fig S2).

We first examined whether CnoIDRΔHΔFAB restored adult viability, by crossing Balanced mutants to animals carrying the null allele, *cno^R2^*/TM3. *cnoIDRΔH/cno^R2^* adult viability parallels that of a wildtype *cno* gene (33% versus 33% expected—Balancer homozygotes die; n=602 (Jensen et al., 2025), and we obtained adult progeny of *cnoIDRΔH/cno^R2^* females crossed to males, revealing maternal-zygotic adult viability. We next assessed viability of *cnoΔFAB/cno^R2^* adults. Viability was substantially reduced (12% versus 33% expected; n=820), though one can generate a homozygous viable stock (Perez-Vale et al., 2021). If the two F-actin binding sites were redundant, we suspected deleting both might lead to adult lethality. To our surprise, *cnoIDRΔHΔFAB/cno^R2^*adults were viable (Fig 6B). However, their viability was even lower than that of either single deletion (8% versus 33% expected; n=681). This is the lowest adult viability of any *cno* mutant we previously tested, other than those that are not adult viable at all. We also could not obtain any adult progeny when we crossed *cnoIDRΔHΔFAB/cno^R2^*females to males, revealing maternal/zygotic lethality. Thus, by this measure, deleting both F-actin binding sites did not eliminate Cno function, but did reduce it relative to deleting either single F-actin binding site.

### Deleting both F-actin binding sites impairs function more than deleting either alone

Next, we used our sensitized assay for function, crossing *cnoIDRΔHΔFAB/cno^R2^*males and females, and comparing the results to our earlier analysis of *cnoΔH* and *cnoΔFAB.* 25% of the progeny are zygotically homozygous for *cno^R2^,* 50% are *cnoIDRΔHΔFAB/cno^R2^*, and 25% are zygotically wildtype. Crosses of *cnoIDRΔH/cno^R2^* flies led to slightly elevated lethality (29%; (Jensen et al., 2025), suggesting CnoIDRΔH provides nearly wildtype function. In contrast, CnoΔFAB retained less function: crosses of *cnoΔFAB/cno^R2^* flies led to 75% embryonic lethality (Perez-Vale et al., 2021). This was further enhanced when both F-actin binding stickers were deleted. 88% of the progeny of *cnoIDRΔHΔFAB/cno^R2^* females and males died (n=266), suggesting that all or almost all *cnoIDRΔHΔFAB/cno^R2^* progeny died. We then examined cuticles of dead embryos. While progeny of both *+/cno^R2^* and *cnoIDRΔH/cno^R2^* largely exhibit only defects in head involution (as in Fig 6D or E; quantified in I), 33% of *cnoIDRΔHΔFAB/cno^R2^* progeny exhibit failure of both head involution and dorsal closure (as in Fig 6G,H; quantified in I). In contrast, failure of both head involution and dorsal closure occurs in only 18% of *cnoΔFAB/cno^R2^* progeny (Perez-Vale et al., 2021). Thus, the phenotypes of *cnoIDRΔHΔFAB* are substantially stronger than those of *cnoIDRΔH*, and modestly more severe than those of *cnoΔFAB*, confirming at least partial redundancy of the two stickers.

### *cnoIDRΔHΔFAB* is the most severe of our viable alleles but less severe than deletion of both spacers

Since *cnoIDRΔHΔFAB/cno^R2^* is adult viable, this protein retains significant function relative to mutants lacking the entire proximal IDR. To make a direct comparison with *cnoΔProxIDR*, *cnoIDRHelixOnly,* or *cnoΔIDR*, we generated maternal-zygotic *cnoIDRΔHΔFAB* mutants. Maternal-zygotic *cno* null mutants all die as embryos with fully penetrant defects in both dorsal closure and head involution, most of which also have defects in epidermal integrity (Fig 6J; (Gurley et al., 2023). Maternal-zygotic *cnoΔProxIDR* mutants also all die as embryos, but their morphogenetic defects are substantially milder than the null mutant, with only 42% exhibiting failure of dorsal closure and few with the strong epidermal integrity defects common in null mutants (Fig 6J; (Jensen et al., 2025). Strikingly, only 34% of maternal-zygotic *cnoIDRΔHΔFAB* mutants died (n=908). Cuticle phenotypes of the dead embryos were overall weak, with most exhibiting only defects in head involution—when we included the maternal-zygotic mutants who hatched 70% of the maternal-zygotic mutants were wildtype in cuticle phenotype (Fig 6J). This places CnoIDRΔHΔFAB protein in a new place in the functional spectrum. It is less functional than any single segment deletion except deleting RA1. However, CnoIDRΔHΔFAB is substantially more functional than either mutant removing both spacers l(CnoΔProxIDR or CnoIDRHelixOnly) as these have fully penetrant maternal/zygotic lethality and much stronger cuticle phenotypes (Fig 6J; (Jensen et al., 2025). While some *cnoIDRΔHΔFAB* maternal/zygotic mutants hatched as larvae, none survived to adulthood (relative to 249 paternally rescued siblings). This contrasts with the maternal/zygotic viability of both *cnoΔFAB/cno^R2^* and *cnoIDRΔH/cno^R2^*(Jensen et al., 2025; Perez-Vale et al., 2021).

These data also revealed a dependence of phenotype severity on gene and likely protein dosage. Maternal/zygotic mutant embryos analyzed receive a full dose of maternal and zygotic mutant protein. In this situation *cnoIDRΔHΔFAB* cuticle phenotypes are quite mild. In contrast, in our analysis above using progeny of *cnoIDRΔHΔFAB/cno^R2^* animals, all progeny have 50% reduction in maternal contribution, and embryos vary from full to 50% to no zygotic contribution. These have more severe cuticle phenotypes. Intriguingly, this was also true in our analysis of *cnoΔFAB*. Progeny of homozygous mutants with full maternal and zygotic contribution exhibited 25% lethality while progeny of *cnoΔFAB/cno^R2^* animals exhibited 75% lethality. suggesting almost all *cnoΔFAB/cno^R2^* progeny died (Perez-Vale et al., 2021).

### CnoIDRΔHΔFAB protein localizes to AJs and retains some TCJ enrichment

Our previous analyses revealed a role for the proximal IDR in effective AJ localization, a role for each spacer in full TCJ enrichment, and a role for sequences in the C spacer in nuclear exclusion. Individually deleting either the FAB or the H stickers did not affect any of these properties, including TCJ enrichment (Jensen et al., 2025; Perez-Vale et al., 2021). We thus examined localization of CnoIDRΔHΔFAB to see if the two stickers play redundant roles. CnoIDRΔHΔFAB localized to AJs from the onset of gastrulation through the end of embryonic morphogenesis (e.g., Fig 7A). Like wildtype Cno (Fig. 1I), CnoIDRΔHΔFAB was strongly enriched in apical AJs (Fig 7B, yellow arrows), did not localize to lateral junctions (Fig 7B, cyan arrows), and did not localize to nuclei (Fig 7b, magenta arrow). CnoIDRΔHΔFAB retained some enrichment to TCJs, but this enrichment was reduced relative to wildtype (1.7 fold vs 2.9 fold for wildtype; Fig 7C vs D, quantified in 6E). Thus, the two stickers play partially redundant roles in TCJ enrichment. However, the spacers are even more important for enrichment in AJs under tension. CnoIDRΔC is only 1.3-fold enriched while in CnoIDRHelixOnly, lacking both spacers but retaining both stickers, TCJ enrichment is essentially eliminated (≤1.1 fold; (Jensen et al., 2025). Thus, the stickers are not essential for AJ localization but do enhance the Cno’s localization in response to tension.

**Fig 7.**
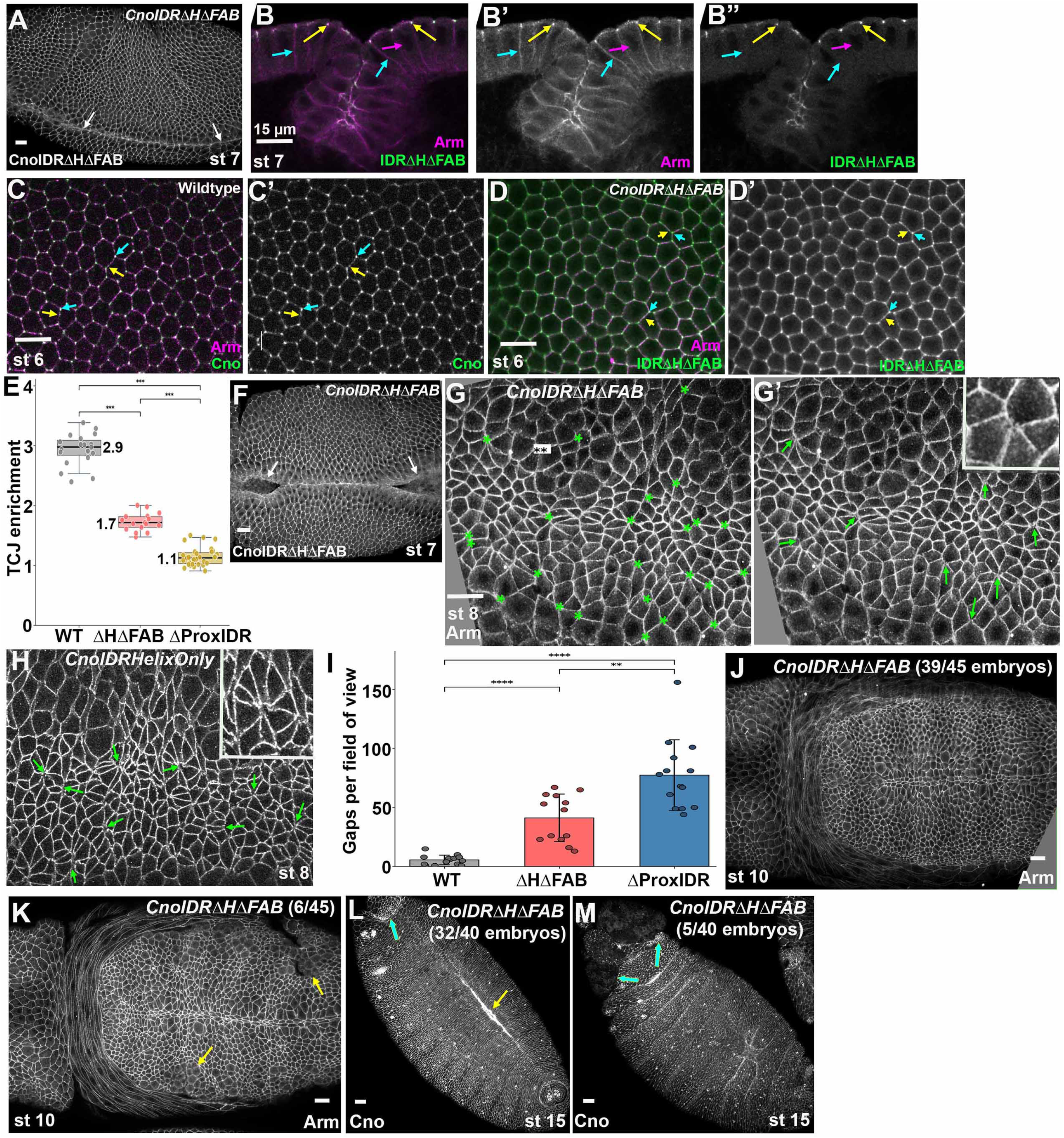
*cnoIDRΔHΔFAB* maternal-zygotic mutants have modest defects in morphogenesis. A-D, F-H, J-M. Embryos, genotypes, embryonic stage, and antigens indicated. Since our Cno antibodies recognize both wildtype Cno and CnoIDRΔHΔFAB, we cannot distinguish maternal/zygotic mutants and paternally-rescued embryos. A. CnoIDRΔHΔFAB localizes to AJs and about half of mutant embryos complete ventral furrow invagination (arrows). B. CnoIDRΔHΔFAB protein localizes to apical AJs and not lateral borders (yellow vs cyan arrows) and does not accumulate in nuclei (magenta arrow). C, D. Wildtype Cno is enriched at TCJs vs bicellular junctions (C, cyan vs yellow arrows). TCJ enrichment of CnoIDRΔHΔFAB is reduced (D, cyan vs yellow arrows). E. Quantification of TCJ enrichment. F. Moderate defects in ventral furrow invagination are seen in about half of mutant embryos. G. Small gaps in AJs in *cnoIDRΔHΔFAB* mutants (e.g inset). H. *cnoIDRHelixOnly* mutants have larger AJ gaps (e.g., inset). I. Quantification of gap frequency. J, K. In most mutants cells rapidly resume columnar shape after mitosis (J), but in a subset this is delayed. L, M. Most embryos complete dorsal closure (L, yellow arrow) and are proceeding with head involution (L, cyan arrow), but in some head involution has failed (M, arrows).

### *cnoIDRΔHΔFAB* mutants have defects in AJ integrity under tension but many morphogenetic movements are completed successfully

The first morphogenetic event requiring Cno is mesoderm invagination. *cno* null mutants have fully penetrant defects (Sawyer et al., 2009). Because CnoIDRΔHΔFAB is recognized by our Cno antibody, we cannot distinguish maternal/zygotic mutants from paternally rescued siblings, so that issue needed to be taken into account in our phenotypic analysis. Assuming all paternally rescued animals invaginate mesoderm correctly, 71% of *cnoIDRΔHΔFAB* maternal-zygotic mutants fail to do so (Table S1; Fig 7A; n=24). This level of defects is substantially elevated relative to *cnoIDRΔ*H mutants (12% failed to invaginate mesoderm correctly) and similar to *cnoΔFAB* homozygotes (67% failed to close normally; (Perez-Vale et al., 2021). *cnoIDRΔHΔFAB* furrow invagination defects ranged from mild to severe (Fig 7F; Table S1). In this parameter, it was only slightly less severe than *cnoΔProxIDR* or *cnoIDRHelixOnly* where 81% or 78% fail to correctly invaginate their mesoderm (Table S1; (Jensen et al., 2025).

Cno reinforces AJs under elevated tension, especially those at aligned anterior-posterior borders or the center of rosettes. Gaps are rare in wildtype (Fig 3K), but in Cno’s absence, gaps form in AJs where tension is highest (Sawyer et al., 2011). We thus assessed gap frequency in *cnoIDRΔHΔFAB* progeny. Many embryos had multiple gaps or disruptions at rosette centers, small bicellular junctions and along aligned AP borders), but most were small disruption (Fig 7G, G’; quantified in 6I; mean 39 gaps/field). Gap frequency was similar to that in *cnoΔProxIDR* mutants (38 gaps/field) but lower than that in *cnoIDRHelixOnly* mutants (77 gaps/field), and gaps in *cnoIDRΔHΔFAB* progeny were smaller than those in *cnoIDRHelixOnly* mutants (Fig 7G vs H).

As cells continue to round up to divide in stages 9-11, cells in the ventral epidermis of *cno* null mutants and of strong mutants like *cnoΔRA* or *cnoIDRHelixOnly* have difficulty resuming columnar architecture and junctions are disrupted (Jensen et al., 2025; Perez-Vale et al., 2021), reflected in the ventral epidermal holes observed in cuticles. In contrast, 87% of stage 9-11 *cnoIDRΔHΔFAB* progeny appeared normal (Fig 7J; 39/45 observed), while 13% had mild delays in cells returning to epithelial architecture (Fig 7K; 6/45 observed). Null and strong *cno* mutants also have penetrant defects in dorsal closure and head involution. The cuticle defects of *cnoIDRΔHΔFAB* progeny suggested its end-stage phenotype would be much weaker, as some mutants hatch and others die with wildtype cuticles (Fig 6J). Consistent with our cuticle analysis, 32/40 stage 15 embryos appeared to complete dorsal closure and head involution correctly (Fig 7L), 3/40 had mild defects in dorsal closure, while in 5/40 head involution failed (Fig 7M). Taken together these phenotypic analyses reveal that the two F-actin binding stickers are somewhat redundant in function, with phenotypes of *cnoIDRΔHΔFAB* embryos consistently worse than those with a deletion of a single sticker. However, CnoIDRΔHΔFAB protein retains a surprising amount of function, substantially more than the mutant missing both spacers.

### *cnoIDRΔHΔFAB* mutants have more severe defects in eye development than either single mutant

As a final test of CnoIDRΔHΔFAB, we examined its function in pupal eye development. Mutants lacking each of the individual F-actin-binding stickers, *cnoIDRΔH* and *cnoΔFAB,* have relatively mild defects. To make direct comparisons with *cnoIDRΔHΔFAB* we compared each mutant heterozygous with the *cno* null allele *cno^R2^*. *cnoIDRΔH/cno^R2^*mutants have few defects (Fig. 4G, J; (Jensen et al., 2025), like wildtype (0.48 versus 0.61 defects per ommatidium, respectively). In *cnoΔFAB/cno^R2^* mutants defect frequency was slightly elevated (0.92 defects per ommatidium; Fig. 4H, J), similar to what we observed with *cnoΔFAB* homozygotes (Perez-Vale et al., 2021). In contrast, *cnoIDRΔHΔFAB/cno^R2^* mutants had a highly elevated frequency of defects (3.98 defects per ommatidium; Fig. 4I, J). This was the highest frequency of defects among all our adult viable mutants. Further, the numerical scoring likely understates severity, as many ommatidia had defects severe enough that the usual scoring system did not capture them all. Thus, in this assay as well, CnoIDRΔHΔFAB protein was significantly less functional than either single sticker deletion. However, it is adult viable, allowing us to assess eye phenotypes, thus contrasting with mutants deleting the entire IDR or deleting both spacers.

## Discussion

One fundamental cell biology discovery of the past decade is the key role played by IDRs in assembly and function of biomolecular condensates. Condensates spatially restrict and concentrate the relevant molecules to enhance function via kinetics and avidity (Holehouse and Kragelund, 2024). Cell-cell and cell matrix adhesion are mediated by large, dynamic protein complexes, and many junctional proteins contain IDRs that play important roles in their function (Rouaud et al., 2020; Sun et al., 2022). However, we are just beginning to dissect the mechanisms by which individual motifs within these IDRs confer protein function. We focus on Cno and its mammalian homolog Afadin, which reinforce connections between cadherin-based junctions and the actomyosin cytoskeleton in response to force. Their IDRs contain conserved “stickers” binding F-actin and other partners, and less well-conserved spacers (Gurley et al., 2023). Here we define the role of these motifs in protein localization and function during morphogenesis.

### Cno’s IDR is essential for its function

Cno and Afadin share five N-terminal folded domains and long C-terminal IDRs (Fig. 1A). Their IDRs diverge dramatically in sequence, length, charge, and amino acid composition (Gurley et al., 2023). Biochemical analyses drew our attention to two conserved stickers, H and the FAB. Both can bind F-actin, and H also stabilizes actin-alpha-catenin interactions (Gong et al., 2025; Jensen et al., 2025). These stickers are separated from the PDZ domain and from one another by poorly conserved spacers, N and C, respectively. Our earlier work revealed a surprising fact: no individual IDR segment is essential for protein function—in fact animals with deletions of H, the FAB, N, or C, are all adult viable and fertile (Fig 8; (Jensen et al., 2025; Perez-Vale et al., 2021). However, sensitized assays revealed that none of these mutant proteins confer fully wildtype function. The C spacer also helps mediate both Cno recruitment to AJs under tension and nuclear exclusion. In contrast, deleting the entire proximal IDR (CnoΔProxIDR), spanning N, H, and C, substantially reduces Cno function, leading to penetrant embryonic lethality (Fig 8; (Jensen et al., 2025). However, CnoΔProxIDR retains residual function. Defects in dorsal closure are less frequent than in *cno* null mutants, disruptions of epidermal integrity are rare and gaps at cell junctions under tension were smaller than those seen in the absence of Cno (Gurley et al., 2023; Jensen et al., 2025). CnoΔProxIDR also had reduced junctional localization, instead localizing to nuclei. A parallel mutant of Afadin (AfadinΔC) also exhibited reduced function, reduced junctional localization and nuclear accumulated in nuclei (Kuno et al., 2025)..

**Figure 8.**
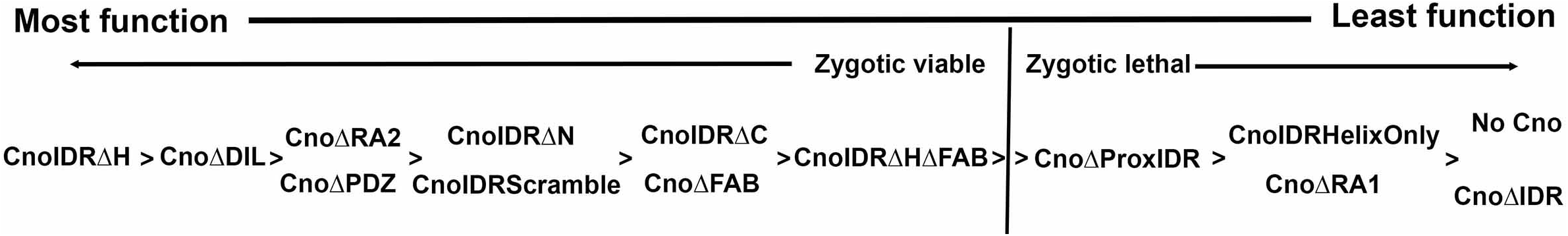
Diagrammatic illustration of the relative severity of different *cno* mutants.

Both CnoΔProxIDR and AfadinΔC retained the FAB, which might mediate their residual function. To test this, we created CnoΔIDR, removing the entire C-terminal IDR but leaving all the folded protein domains intact. Both CnoΔIDR and CnoΔProxIDR have reduced AJ localization and instead accumulate in nuclei. Both are no longer enriched at AJ under tension and accumulate at the cortex of mitotic cells after being released from nuclei. Since CnoΔIDR retains weak AJ recruitment, the folded protein domains alone can mediate some enrichment there. However, the embryonic phenotype of *cnoΔIDR* is much stronger than that of *cnoΔProxIDR*, with penetrant disruption of dorsal closure, frequent disruption of ventral epidermal integrity and larger gaps at cell junctions under tension. Thus, the IDR is important for Cno localization and is essential for Cno function (Fig 8). The presence of the FAB in CnoΔProxIDR clearly provides some residual function--we return to the potential redundancy of the two F-actin binding stickers below. Future work should define how which, if any, functional deficits of CnoΔIDR and CnoΔProxIDR are due to nuclear sequestration, by reducing AJ localization. Nuclear localization alone does not fully abrogate function, as both CnoIDRΔC and CnoIDRScramble localize to nuclei, but this could be a matter of degree. Constructs tethering CnoΔIDR to the plasma membrane, preventing nuclear import, could test this.

### IDR spacer length is more important than sequence, but the C spacer has embedded sequences mediating nuclear exclusion

While mutants individually deleting the N and C spacers are viable, deleting both spacers while retaining the F-actin binding stickers (CnoIDRHelixOnly) strongly reduces Cno function, leading to embryonic lethality and strong defects in morphogenesis (Fig 8; (Jensen et al., 2025). CnoIDRHelixOnly also has strongly reduced AJ localization and instead re-localizes to nuclei. To test whether space sequences were critical, or whether length and amino acid composition were more important, we scrambled both spacer sequences. *cnoIDRScramble* mutants were viable and fertile, and CnoIDRScramble protein accumulated in apical AJs. Thus, while the presence of the spacers is critical, spacer sequence is not essential. Our sensitized assays in embryos and pupal eyes suggest that *cnoIDRScramble* mutant phenotypes overall are slightly stronger than those of *cnoIDRΔN* mutants, and similar to or slightly weaker than those of *cnoIDRΔC* mutants (Fig 8). This is consistent with the idea that the C spacer has important sequences embedded within it which are disrupted by scrambling. These likely include sequences mediating nuclear exclusion and localization to AJs under tension, as CnoIDRScramble protein shares with CnoIDRΔC protein significant deficits in these properties. Cno and its Dipteran homologs share a consensus sequence for binding the nuclear export factor Exportin in spacer C (LAKELNQLTM in Cno, matching the φ-x₂₋₃-φ-x₂₋₃-φ-x-φ consensus; (Xu et al., 2012). Replacing Cno’s proximal IDR with that of Afadin, which shares no detectable sequence similarity to Cno’s IDR outside of the conserved helices and no apparent match to the Exportin consensus, created a viable mutant with reduced function, elevated nuclear localization and reduced enrichment at TCJs similar to that of CnoIDRΔC (Jensen et al., 2025).

### The F-actin binding stickers are somewhat redundant in function but are not essential for viability

While Cno and Afadin share long C-terminal IDRs, there are only two sequences with detectable conservation: the predicted helices in the center of the IDR (the H sticker) and those at the C-terminus (the FAB (Gurley et al., 2023). Both can bind F-actin, and the H also stabilizes actin:alpha-catenin interactions (Gong et al., 2025; Jensen et al., 2025). We used AlphaFold to model the FAB interaction (Jensen et al., 2025). Here we probed this model by mutating conserved phenylalanines predicted to be involved in F-actin binding: this reduced F-actin binding, further supporting this model. Afadin’s proximal IDR can largely replace Cno’s proximal IDR in our functional assays, even though the H segment and the FAB are the only conserved regions, initially suggesting the hypothesis that each of these two conserved segments might be essential. However, mutants individual deleting either the FAB or H are viable (Jensen et al., 2025; Perez-Vale et al., 2021)—in fact *cnoIDRΔH* mutants are the least affected of all the Cno segmental deletions we assessed.

We thus tested the hypothesis that the relatively mild functional deficits of CnoΔFAB and CnoIDRΔH are because the two F-actin binding segments are redundant in function, by generating *cnoIDRΔHΔFAB.* If at least one F-actin binding site was essential, its function would be strongly or completely disrupted, as we saw when we deleted both spacers. Instead, the results were more complex. In most assays, *cnoIDRΔHΔFAB* mutant phenotypes were more severe than those of individual deletions (*cnoΔFAB* and *cnoIDRΔ*H; Fig 6). Most striking, zygotic adult viability of *cnoIDRΔHΔFAB* mutants was strongly reduced to levels lower than those of either individual deletion, and, unlike *cnoΔFAB* and *cnoIDRΔH,* maternal/zygotic mutants were not viable to adulthood. However, CnoIDRΔHΔFAB protein was substantially more functional than CnoΔProxIDR or CnoIDRHelixOnly. The latter mutants are zygotic lethal. Further, while *cnoΔProxIDR* or *cnoIDRHelixOnly* maternal/zygotic mutants die as embryos with strong defects in morphogenesis, some *cnoIDRΔHΔFAB* maternal/zygotic mutants survive embryogenesis and most that die only have modest defects (Fig 8). Thus, while the two conserved F-actin binding stickers are somewhat redundant in function, our results invalidate our initial hypothesis that they would be the most essential parts of Cno’s IDR. In fact, they are less essential than the two non-conserved spacers, as simultaneously deleting both spacers leads to much stronger defects. Since the stickers are the only regions of Cno/Afadin known to bind F-actin, this also suggests that either there is an additional F-actin binding site, or Cno/Afadin enhances AJ:cytoskeletal linkage more indirectly.

Our analysis of *cnoIDRΔHΔFAB* also revealed another surprise. While *cnoIDRΔHΔFAB* maternal/zygotic mutants have only mild defects in embryonic morphogenesis and some hatch, embryos in which the levels of maternal and zygotic CnoIDRΔHΔFAB protein are reduced (as in progeny of *cnoIDRΔHΔFAB/cno^R2^* flies) have more severe morphogenetic defects. We saw similar differences when comparing *cnoΔFAB* maternal/zygotic mutants and the progeny of *cnoΔFAB/cno^R2^* flies (Perez-Vale et al., 2021). Thus, when the function of some mutant Cno proteins are reduced, their levels become important. It is tempting to speculate that this might be related to threshold effects in assembling protein complexes by multivalent interactions.

Stepping back, what did we learn about the function of Cno’s IDR and its mechanism of action? Broadly, the Cno IDR is essential for protein function but it is very robust to mutation. No individual segment is essential, including the only two segments conserved between flies and mammals. Scrambling the sequences of the two spacers reduced but did not eliminate function, suggesting their length and amino acid composition are more important than their sequence. Surprisingly, the spacers were more important than the stickers, as is illustrated both by individual or dual deletions. This raises questions for the future. Does the IDR play a specific role in bridging AJs to the F-actin network across a requisite distance? Are there yet to be mapped cryptic stickers in the N and C spacers, potentially including a third F-actin binding site, or do the spacers purely serve entropic functions?

We suspect there will be similarities but few universal roles governing IDR function. Contrasting Cno with some select examples illustrates this. Analysis of the C-terminal IDR of the transmembrane protein Linker for Activation of T cells (LAT) revealed a small number of sequence motifs whose alteration severely affected function (Rubin et al., 2026). The authors’ powerful ability to assess subtle quantitative effects also revealed many more segments with moderate or mild effects on function. Thus, in LAT’s IDR, multiple individual segments act in a combinatorial fashion. The long C-terminal IDR of Abelson kinase (Abl) provides a contrasting example (Cheong et al., 2020; Cheong and VanBerkum, 2017; Duan et al., 2023; Rogers et al., 2021; Rogers et al., 2016). It has multiple segments conserved in insects or vertebrates, but only one, an SH3-binding motif, is conserved between flies and mammals. Only this motif is critical for function, and all other segments can be individually deleted, including the C-terminal F-actin binding site. However, deleting the entire IDR has more drastic effects than any individual deletion, as we observed with Cno. The APC protein, part of the Wnt-regulatory destruction complex, provides a third example. It’s IDR contains multiple stickers conserved between flies and mammals, flanked by poorly conserved spacers (Stamos and Weis, 2013). These stickers include multiple copies of two motifs binding beta-catenin, multiple copies of a motif binding the destruction complex partner Axin, and a final conserved sequence without a known binding partner. Once again, many individual motifs are dispensable—function appears to require one beta-catenin binding site, one Axin binding site, and the other conserved sequence (Kunttas-Tatli et al., 2015; Kunttas-Tatli et al., 2012; Roberts et al., 2011; Smits et al., 1999). Deleting the entire IDR essentially eliminates function, and in fact mutations in colorectal tumors select for partial loss-of-function (Shibata et al., 1997; Smits et al., 1999). Thus, in APC’s IDR, multiple stickers are important, but as most are present in multiple copies, redundancy is prominent. These four examples provide a glimpse of the complexity of the “sequence based grammar” that determines IDR specificity and function. To develop predictive models will require combining detailed individual experimental datasets like ours with new computational approaches as machine learning (Chong and Mir, 2021; Holehouse and Alberti, 2025).

## Acknowledgements

We are very grateful to Dr. Nat Prunet of the Biology Imaging Core for imaging advice and support, Rachel Szymanski for help getting this project off the ground, Noah Gurley for advice on immunoblotting, Tara Finegan, Scott Williams and Bob Duronio for comments on the manuscript, and Peifer, Bergstralh, Finegan, Lovegrove, and Williams lab members for helpful discussions. This work was funded by NIH R35 GM118096 to M. Peifer. The authors declare no competing financial interests.

## Author contributions

Corbin Jensen and Renick Wiltshire led analysis of protein localization and embryonic phenotypes. Sarah Clark analyzed pupal eye phenotypes. Jeremy Lamb carried out immunoblotting analysis. Kevin Slep carried out the biochemical analyses. Mark Peifer helped with fly genetics. The manuscript was written by Mark Peifer and Kevin Slep, with editorial input from all of the authors.

## Methods

### Drosophila work

All experiments were performed at 25°C unless noted otherwise. we used flies of the *yellow white* genotype [Bloomington Drosophila Stock Center (BDSC), stock 1495] as controls and they are referred to in the text as wildtype (WT). Maternal/zygotic mutants of *cnoΔIDR* and *cnoIDRΔHΔFAB* were made using the FRT/ovoD approach. We heat-shocked third-instar larvae generated by crossing *cno*Δ*IDR/TM3* or *cnoIDRΔHΔFAB /TM3* virgin females to *P{ry[+t7.2]=hsFLP}1, y[1] w[1118]; P{neoFRT}82B P{ovo−D1−18}3R/TM3, ry[*], Sb[1]* males in a 37°C water bath for 2 hours each on two consecutive days. Virgin female adult progeny with the germline genotype *hsFLP1; P{neoFRT}82B P{ovo−D1−18}3R/P {neoFRT}82B cnoΔIDR* (or *cnoIDRΔHΔFAB)* were collected and subsequently crossed with *cnoΔIDR* (or *cnoIDRΔHΔFAB) /TM3 Sb* males. Embryos generated from this cross were analyzed. For *cnoΔIDR* crosses maternal/zygotic mutants identified by absence of staining with our Cno antibody which recognizes an epitope in the proximal IDR, and referred to in the text as simply *cnoΔIDR*. Sensitized assays were done using mutant flies that are heterozygous for each *cno-mutant* allele with the null allele *cno^R2^* as described in (Perez-Vale et al., 2021).

### Cloning of cno IDR structure-function constructs

Most constructs were generated using Azenta Life Science (Waltham, MA, USA) cloning and mutagenesis service. Details and sequences are in Fig S1. *cnoRA-PDZ* was generated using *cnoΔProxIDR* as an initial template (Jensen et al., 2025), deleting all nucleotides corresponding to Canoe amino acids Y1927-H2051. *cnoIDRΔHΔFAB* was generated using *cnoIDRΔH* as an initial template (Jensen et al., 2025), deleting all nucleotides corresponding to Canoe amino acids Q1937-H2051. *cnoIDRScramble* was designed using the Shuffle Protein server (https://www.bioinformatics.org/sms2/shuffle_protein.html)(Stothard, 2000) as follows: the Canoe IDR-N region (residues G1130-Y1565 as well as the preceding glycine from the PAGG cloning site insert) and the Canoe IDR-C region (residues A1671-Q1926) were independently entered into to Shuffle Protein, which generated the respective scrambled sequences shown in Fig. S1B. The scrambled sequences were verified for predicted disorder using the Metapredict server (https://metapredict.net/) (Lotthammer et al., 2026), DNA encoding the full proximal IDR [IDR-N Scramble - H region – IDR-C Scramble] was codon-optimized for Drosophila melanogaster expression, flanked by SbfI and SpeI sites, and synthesized by GeneArt (Thermo Fisher Scientific, Waltham, MA, USA). The [IDR-N Scramble - H region – IDR-C Scramble] region was then inserted into the SbfI and SpeI sites of *cnoCanoeIDR* by Azenta Life Science, replacing the Canoe IDR to generate *cnoIDRScramble*.

All constructs were verified by whole plasmid DNA sequencing (Plasmidsaurus, South San Francisco, CA, USA). Each construct contains the *w^+^* selectable marker and is flanked by attR and attL sites and was integrated into the attP site at the *cnoΔΔ* locus (Bloomington Drosophila Stock Center stock 94023; (Perez-Vale et al., 2021). DNA was injected into *PhiC31/intDM. Vas; cnoΔΔ* embryos (BDSC stock 94023) by BestGene (Chino Hills, CA, USA). F1 offspring were screened for the presence of the *w^+^* marker and outcrossed to *w; TM6B, Tb/TM3, Sb* (BDSC stock 2537) to generate a balanced stock over TM3. Each stock was verified by PCR, sequencing and western blot analysis. We outcrossed these stocks to a *y w* stock with a wild type 3rd chromosome to remove potential passenger mutations from the mutant chromosomes. This occurred for multiple generations selecting for the linked *w^+^* marker in each generation. We were thus able to generate homozygous stocks for *cnoIDRScramble*.

### Immunoblotting and quantification

To determine the relative levels of wildtype Cno, GFP-tagged mutants, and α-tubulin we used immunoblotting of embryonic lysates collected at 4-8 hours after egg laying. We generated embryonic lysates were generated as in (McParland et al., 2024b). Embryos were collected into 0.1% Triton X-100, dechorionated in 50% bleach for 5 minutes, and washed three times with 0.1% Triton X-100. Next, lysis buffer (1% NP-40, 0.5% Na deoxycholate, 0.1% SDS, 50 mM Tris pH 8.0, 300 mM NaCl, 1.0 mM DTT, HaltTM Protease and Phosphatase Inhibitor Cocktail (Thermo Fisher Scientific, #78442) (100×), and 1 mM EDTA) was added. Embryos were then ground with a pestle for ∼20 seconds and then placed on ice for 10 minutes. Embryos were then ground again with a pestle for ∼20 seconds and subsequently centrifugated at 16,361xg for 15 minutes at 4°C. We assessed protein concentration using the Bio-Rad Protein Assay Dye, recording absorbance at 595 nm with a spectrophotometer. We then ran protein lysates on 7% SDS-PAGE and transferred proteins onto nitrocellulose membranes. The membranes were blocked with 5% bovine serum albumin (BSA) diluted in Tris-buffered saline with 0.1% Tween-20 (TBST) for 1 hour at room temperature. Primary and secondary antibodies were diluted in 5% BSA diluted in TBST. Primary antibody incubations were done overnight at 4°C, and secondary antibody incubation for 1 hour at room temperature. We used the Odyssey CLx infrared system (LI-COR Biosciences) to image the membranes, and band densitometry was carried out using Empiria Studio® Software (LI-COR Biosciences).

### Embryo Fixation and Immunofluorescence

Fixation and immunofluorescence were performed as described previously (Jensen et al., 2025). Briefly; adult flies were transferred to collection cups fitted with apple juice agar plates supplemented with yeast paste. Embryos were harvested with a paint brush in 0.1% Triton-X-100 and subsequently dechorionated by treatment with 50% bleach, followed by three washes in 0.03% Triton X-100 / 68 mM NaCl. Heat fixation was carried out by immersing embryos in a solution of 0.03% Triton X-100, 68 mM NaCl, and 8 mM EGTA at 95°C for 10 seconds, after which samples were transferred to ice and cooled for 30 minutes. Devitellinization was achieved by vigorous agitation in a 1:1 mixture of n-heptane and 95% methanol:5% EGTA, followed by three rinses in 95% methanol / 5% EGTA. For immunostaining, embryos were rinsed three times in phosphate-buffered saline (PBS) supplemented with 5% normal goat serum (NGS; Thermo Fisher Scientific) and 0.1% saponin (PBSS-NGS), then blocked for one hour in PBSS-NGS containing 1% NGS. Primary antibodies were diluted in PBS containing 1% bovine serum albumin and 0.1% saponin (see antibody dilution table) and applied overnight at 4°C. Embryos were subsequently washed three times in PBSS-NGS (15 minutes per wash) and incubated with secondary antibodies for 2 hours at room temperature. Samples were mounted on glass slides in a homemade Gelvatol mounting medium (protocol courtesy of the University of Pittsburgh Center for Biological Imaging). Imaging was performed on a LSM 880 confocal laser-scanning microscope (Carl Zeiss, Jena, Germany) using a 40X/NA 1.3 Plan-Apochromat oil-immersion objective and ZEN Black 2009 acquisition software. Post-acquisition, all images were uniformly adjusted for input levels, brightness, and contrast in Photoshop (Adobe, San Jose, CA).

### Eye disc fixation, imaging and scoring

Prepupae were collected from crosses maintained at 25℃. The samples were stored in humidified chambers and aged until 40 hours after puparium formation (APF). At 40 APF, pupae were placed in a droplet of 1.5x PBS. The fly was removed from the pupal casing, and decapitated. The head casing was then removed, exposing the brain complex. The brain complex was washed in 1.5x PBS twice to remove any excess fat and tissue. The sample was then fixed in 3.7% formaldehyde for 20 min. Following fixation, it was washed in 1.5x PBS twice and blocked in 1.5x PBST for 30 min. The tissue was then placed in primary antibodies overnight at 4℃ (rat anti E-Cadherin (1:50; DSHB) and rat anti N-Cadherin (1:50; DSHB)). After two washes in 1.5x PBS and blocking in 1.5x PBST for 30 min, the tissue was then stained with Goat anti-Rat 568 secondary (1:1000; Life Tech) to detect AJs and retinas. The tissue was imaged with a Zeiss LSM 980 with Airyscan 2 using a 63x/1.4 oil Plan Apochromat objective. Patterning errors were scored in 10-14 eye discs per genotype spanning 88-137 ommatidia per genotype. Data were analyzed for statistical significance using PRISM. Significance was calculated using an ordinary one-way parametric ANOVA. Images were processed for publication using FIJI and Adobe Photoshop.

Prepupae were collected from crosses maintained at 25℃. The samples were stored in humidified chambers and aged until 40 hours after puparium formation (APF). At 40 APF, pupae were placed in a droplet of 1.5x PBS. The fly was removed from the pupal casing, and decapitated. The head casing was then removed, exposing the brain complex. The brain complex was washed in 1.5x PBS twice to remove any excess fat and tissue. The sample was then fixed in 3.7% formaldehyde for 20 min. Following fixation, it was washed in 1.5x PBS twice and blocked in 1.5x PBST for 30 min. The tissue was then placed in primary antibodies overnight at 4℃ (rat anti E-Cadherin (1:50; DSHB) and rat anti N-Cadherin (1:50; DSHB)). After two washes in 1.5x PBS and blocking in 1.5x PBST for 30 min, the tissue was then stained with Goat anti-Rat 568 secondary (1:1000; Life Tech) for 2h at room temperature. The tissue was imaged with a Zeiss LSM 980 with Airyscan 2 using a 63x/1.4 oil Plan Apochromat objective to visualize E-cadherin and N-cadherin in one channel and GFP-tagged Cno proteins in another. Patterning errors were scored in 10-14 eye discs per genotype spanning 88-137 ommatidia per genotype. Data were analyzed for statistical significance using PRISM. Significance was calculated using an ordinary one-way parametric ANOVA. Images were processed for publication using FIJI and Adobe Photoshop.

### Automated TCJ/multicellular junction enrichment analysis

We performed membrane segmentation using the EpySeg graphical user interface (GUI) (Aigouy et al., 2020), which employs a pre-trained deep learning model for image segmentation. Specifically, we utilized the “v2” model from the segmentation_models library (https://github.com/qubvel/segmentation_models), with the following architectural specifications: Model Architecture: Linknet, Backbone Network: VGG16, Activation Function: Sigmoid, and Input Image Dimensions: Dynamically adjusted (0 width and height). The segmentation was performed on representative images, with predictions generated based on the Armadillo staining channel of de-identified images to achieve a single-blinded analysis. Following segmentation, junction detection and intensity quantification were performed using a custom automated Python pipeline implemented across four analysis modules and executed through a PyQt6 graphical user interface. All image processing utilized NumPy, SciPy, scikit-image, tifffile, pandas, and matplotlib libraries.

Tri-cellular junction (TCJ) vertices were identified from the binary skeleton output of EpySeg by detecting branch-point pixels, defined as skeleton pixels with three or more 8-connected skeleton neighbors. Contiguous clusters of branch-point pixels were labeled using connected-component analysis (scipy.ndimage.label), and the centroid of each cluster was taken as the junction coordinate. To ensure measurement quality, junctions whose centroids were within 20 pixels of another junction were excluded, as the intervening bi-cellular branch would be too short to yield a reliable independent measurement. Prior to junction detection, skeletons were re-thinned using morphological skeletonization (skimage.morphology.skeletonize) to enforce a true one-pixel-wide medial axis and eliminate spurious branch points introduced by occasional two-pixel-wide regions in the EpySeg output. Bi-cellular branch lines were traced from each junction cluster outward along non-junction skeleton pixels using a breadth-first search (BFS) algorithm. BFS propagation halted upon reaching any other junction cluster, ensuring each branch was traced in its entirety exactly once and represented a single continuous segment between two junction vertices. Branches whose pixels contacted any border of the image were excluded, and all branches associated with those junctions were similarly removed to prevent truncated edge segments from biasing measurements. Fluorescence intensity in the GFP channel was measured at each tri-cellular junction and along each of its associated bi-cellular branches. TCJ intensity was defined as the mean GFP signal within a circular region of radius 1 pixel centered on the junction centroid. Branch intensity was defined as the mean GFP signal along all branch pixels located more than 5 pixels from the junction centroid, providing a buffer zone that excluded transitional signal at the junction periphery. Branches with fewer than 3 usable pixels after exclusion of the buffer zone were omitted. The TCJ/BCJ enrichment ratio was calculated as the mean TCJ intensity divided by the mean intensity across all valid branches associated with that junction. Per-image summary statistics, including mean TCJ intensity, mean branch intensity, and mean TCJ/BCJ ratio, were computed across all qualifying junctions within each image.

### Analysis of junctional gaps

We selected closeups of similar magnifications of stage 7 and stage 8 embryos of each genotype, stained to visualize Arm which outlines AJs, utilizing lateral views from the ventral midline toward the amnioserosa. These were then randomized, and scored blind for gaps in tricellular or multicellular junctions, or places where the junctional protein signal was broadened. Samples were then unblinded, and total gap number was then recorded per field of view.

### Statistics and quantification

All statistical analyses and figure generation for Tri-cellular enrichment and gap analysis were performed using custom Python scripts. For cross-condition comparisons, the mean TCJ/BCJ intensity ratio for each individual image was used as the unit of biological replication, yielding one value per image. This approach was chosen to avoid inflating statistical power through pseudoreplication of individual junction measurements within an image. Box and whisker plots display the distribution of per-image mean ratios, with boxes representing the interquartile range (25th–75th percentile), the central line representing the median, and whiskers extending to the minimum and maximum values. Individual data points representing each image mean are overlaid on the corresponding box. The grand mean across images is indicated numerically below the lower whisker. Differences across experimental conditions were assessed by one-way ANOVA followed by pairwise independent-samples t-tests with Bonferroni correction for multiple comparisons. Significance thresholds were p < 0.05 (\**), p < 0.01 (**), and p < 0.001 (\*\*\**). Significant pairwise comparisons are indicated by brackets above the plot. Data distribution was assumed to be normal but this was not formally tested.

### Cuticle preparation and analysis

We prepared embryonic cuticles according to (Wieschaus and Nüsslein-Volhard, 1986). Females and males were placed in cups and eggs were collected on apple juice agar plates with yeast for < 24 hours at 25°C. We then placed eggs in groups of ten on a fresh apple juice agar plate without yeast and incubated them at 25°C for 48 hours to allow embryos to develop fully and viable embryos to hatch. We collected unhatched embryos in 50% bleach and dechorionated them for 5 minutes. Bleach was removed and embryos were transferred to glass slides into 100µl of 1:1 Hoyer’s medium:lactic acid, and covered with a cover slip. Slides were incubated at 60°C for 24-48 hours and stored at room temperature. Slides were then imaged, placing each embryo into categories based on morphological criteria, and counting unfertilized eggs and removing them from the total. At least two separate crosses were used for each genotype.

### Cloning and purification of Canoe FAB domain constructs

DNA encoding the wild type *Drosophila melanogaster* Canoe FAB region (residues 1976-2051) was generated using PCR (FAB forward primer: 5’-CGCGGCAGCGGATCCtccactccaggcgtcattggagctcaagaag -3’; FAB reverse primer: 5’-CAAGCTTGTCGACGGAGCTCGAATTCttagtgcaccgcgtctatatctcgttg -3’) while DNA encoding the F2017E/F2024E double mutant was generated by PCR sewing using the above primers along with mutation-encoding primers (mutant forward primer, 5’-tcgGAAaaggagaagatgaagatgGAAgccctggagtcgggggaagataag-3’; mutant reverse primer: 5’-ggcTTCcatcttcatcttctccttTTCcgacagcttttcgggcaccgcatc-3’). DNA was sub-cloned into pGEX-6P2 (Cytiva, Marlborough, MA) using BamH1 and EcoR1 restriction enzymes and T4 DNA ligase (New England Biolabs, Ipswich, MA). All plasmids were sequence verified.

The GST-FAB constructs (as well as a pGEX-6P2 GST alone control construct) were individually transformed into *Escherichia coli* BL21 DE3 pLysS cells and 3 L of each respective culture grown to an optical density at 600 nm of 0.8 in LB broth containing 50 mg/L ampicillin at 37°C. The temperature was lowered to 20°C and protein expression was induced with 100 µM IPTG for 16 h. Cells were harvested by centrifugation, resuspended in lysis buffer (25 mM Tris-HCl pH 8.5, 300 mM NaCl, and 0.1% β-mercaptoethanol (β-ME)) supplemented with 5 μg/ml DNase (Worthington Biochemical Corp., Lakewood, NJ) and 10 μg/ml lysozyme (Thermo Fisher Scientific, Waltham, MA) at 4°C and lysed by sonication. Phenylmethylsulfonyl fluoride was added to 1 mM final concentration. Cells debris was pelleted by centrifugation at 17,000 x g for 45 min. Supernatant was loaded onto a 5 ml Glutathione Sepharose 4 Fast Flow column (Cytiva, Marlborough, MA). The column was washed with 1 L of wash buffer (25 mM Tris-HCl pH 8.5, 300 mM NaCl, and 0.1% β-ME) and the protein batch eluted with 50 mL wash buffer supplemented with 50 mM Glutathione. Eluate was then exchanged into 25 mM Tris-HCl pH 8.5, 300 mM NaCl, and 0.1% β-ME and concentrated using Thermo Fisher Scientific (Waltham, MA) Pierce 10k MWCO centrifugal concentrators (final concentrations were ≥ 190 μM), aliquoted, and stored at −80°C.

### F-actin co-sedimentation assays

F-actin co-sedimentation assays were adapted from (Gong et al., 2025) as follows: Lyophilized rabbit skeletal muscle actin (1 mg; Cytoskeleton, Inc., Denver, CO) was reconstituted to 10 mg/ml (233 μM) with 100 μL in Actin Buffer (2 mM Tris pH 8.0, 0.1 mM CaCl_2_, 0.2 mM ATP, 0.5 mM DTT) and stored at -80°C in 10 μL aliquots. Actin was polymerized by thawing and further diluting to 20 μM by adding 106.5 μl Actin Buffer and mixing with 13 μL 10x KMEI buffer (10X = 500 mM KCl, 10 mM MgCl_2_, 10 mM EGTA, 100 mM imidazole pH 7.0) supplemented with 5 mM ATP and incubated at room temperature for 1 hour (final Actin concentration of 18 μM). Canoe GST-FAB constructs, and the GST control, were diluted in 1X KMEI buffer to a final concentration of 13.5 μM and pre-cleared by ultracentrifugation at 80,000 rpm (278,000 x g) in a TLA-100 rotor (Beckman Coulter, Brea, CA) for 15 minutes at 4°C. 31.1 μL of the supernatant was mixed with either 38.9 μL of 18 μM polymerized actin or 38.9 μL of 1X KMEI buffer to give final concentrations of Canoe at 6 μM and actin at 10 μM. Mixtures were incubated for 30 minutes at room temperature, then ultracentrifuged at 80,000 rpm (278,000 x g) for 30 minutes at 4°C. ¾ of the supernatant (52.5 μL) was mixed with 17.5 μL of 4x SDS-PAGE loading buffer. The remaining ¼ of the supernatant was removed, the pellet was washed twice with 1X KMEI buffer, and then dissolved in 4/3 volume of 1x SDS-PAGE loading buffer (93.3 μL). Samples were separated by SDS-PAGE, stained with Coomassie Brilliant Blue, scanned on a LI-COR Odyssey scanner (LI-COR Biosciences, Lincoln, NE), and quantified using FIJI. Experiments were conducted in quadruplicate. Data were plotted using Prism (GraphPad, San Diego, CA), and analyzed using 2-way ANOVA followed by either Sidak’s or Tukey’s multiple comparisons test.

## Figure Legends

**Fig. S1.**
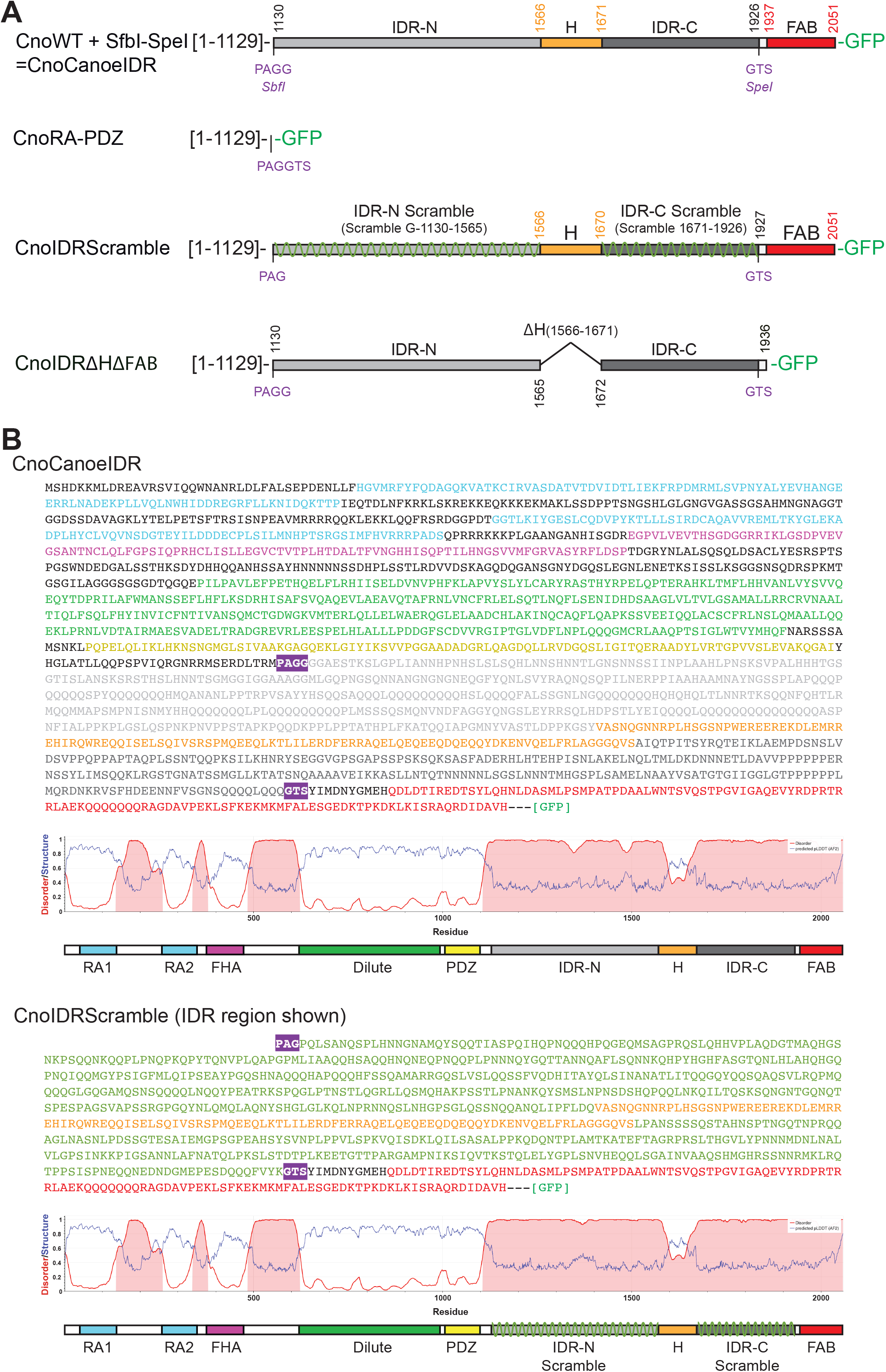
Construct diagrams of Canoe IDR structure-function constructs. A) Domain diagrams of the IDR regions for CnoRA-PDZ, CnoIDRScramble, and CnoIDRΔHΔFAB proteins. The domain diagram of CnoCanoeIDR {Jensen, 2025 #6331}, which served as a template for subsequent constructs, is shown at top. The engineered SbfI and SpeI restriction enzyme (RE) sites, and corresponding amino acid residues are shown in purple. Cloning artifacts in constructs derived from these RE sites are also indicated in purple. All constructs included a C-terminal GFP fusion. B) Top: sequence of CnoCanoeIDR (full length), color-coded according to the domain diagram shown below the sequence. The Metapredict {Lotthammer, 2026 #6402} plot of the per-residue predicted disorder (red line) and the AlphaFold2 {Jumper, 2021 #5829}. pLDDT values (blue line) are shown. Bottom: sequence of CnoIDRScramble (only showing the IDR region), color-coded according to the domain diagram shown below the sequence. The Metapredict plot of the per-residue predicted disorder (red line) and the AlphaFold2 pLDDT values (blue line) are shown.

**Fig. S2.**
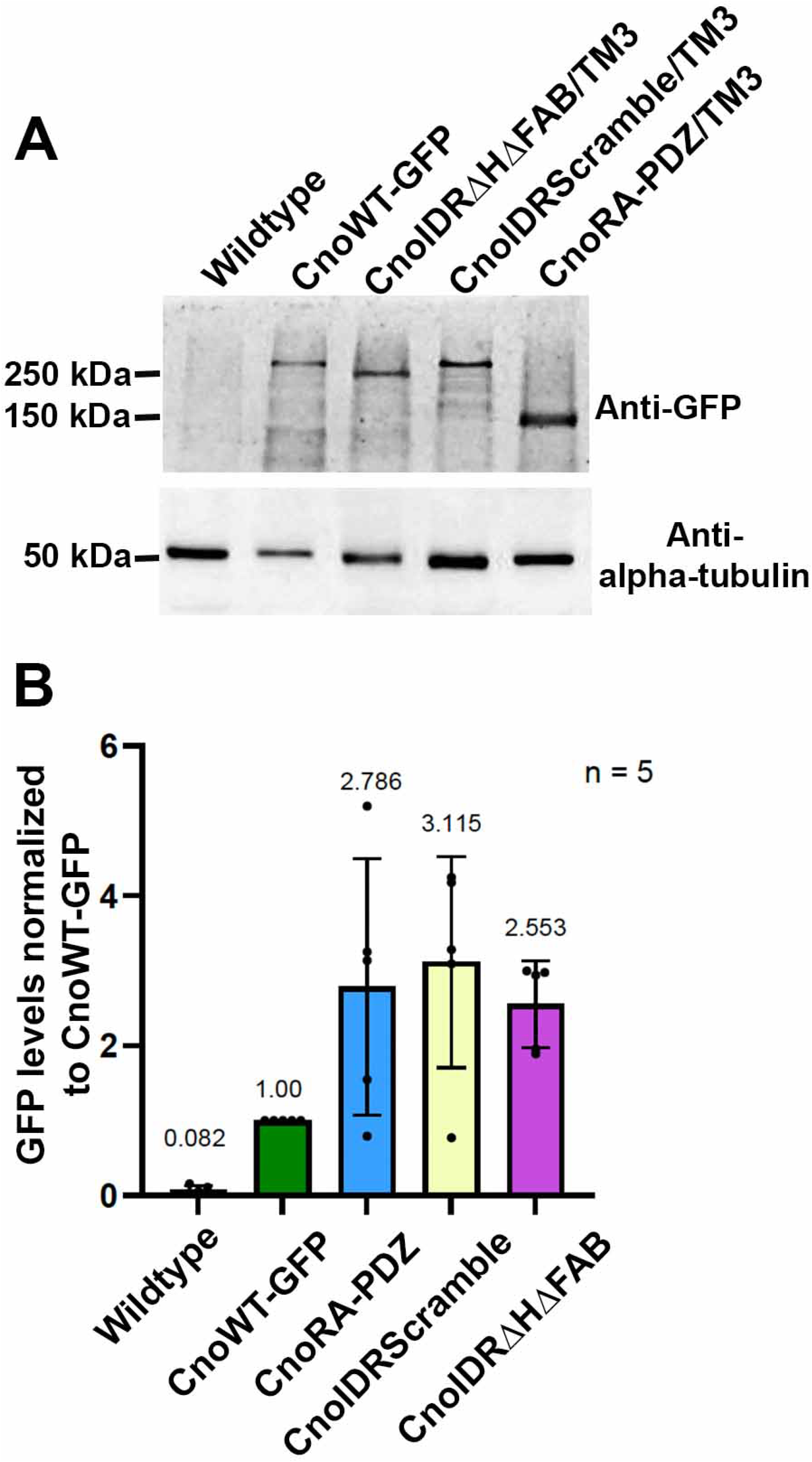
Cno mutant protein accumulation. **(A).** Representative image of an immunoblot from embryos of the indicated genotypes, using protein extracts from 4-8 hour old embryos. Following SDS-PAGE, proteins were transferred onto nitrocellulose membranes and immunoblotted with antibodies against GFP and α-tubulin as a loading control. *y w* (our wildtype control) and homozygous *cnoWT-GFP* embryos served as negative and positive controls for the GFP antibody, respectively. The other mutant samples are from parents heterozygous for the mutant allele and a Balancer chromosome wildtype for *cno*. The protein ladders and the corresponding molecular weights are indicated. **(B).** Quantification of expression levels of Cno mutant proteins relative to CnoWT-GFP, the levels of which were normalized to 1. Since mutant protein extracts were generated from heterozygous animals, expression values for mutant protein levels were multiplied by two as a proxy for homozygous expression, and these values are represented in the graphs. Dots represent replicates –all included at least two biological replicates. Top line of column depicts the mean value, and narrower bands indicate standard error of the mean.

**Table S1.**
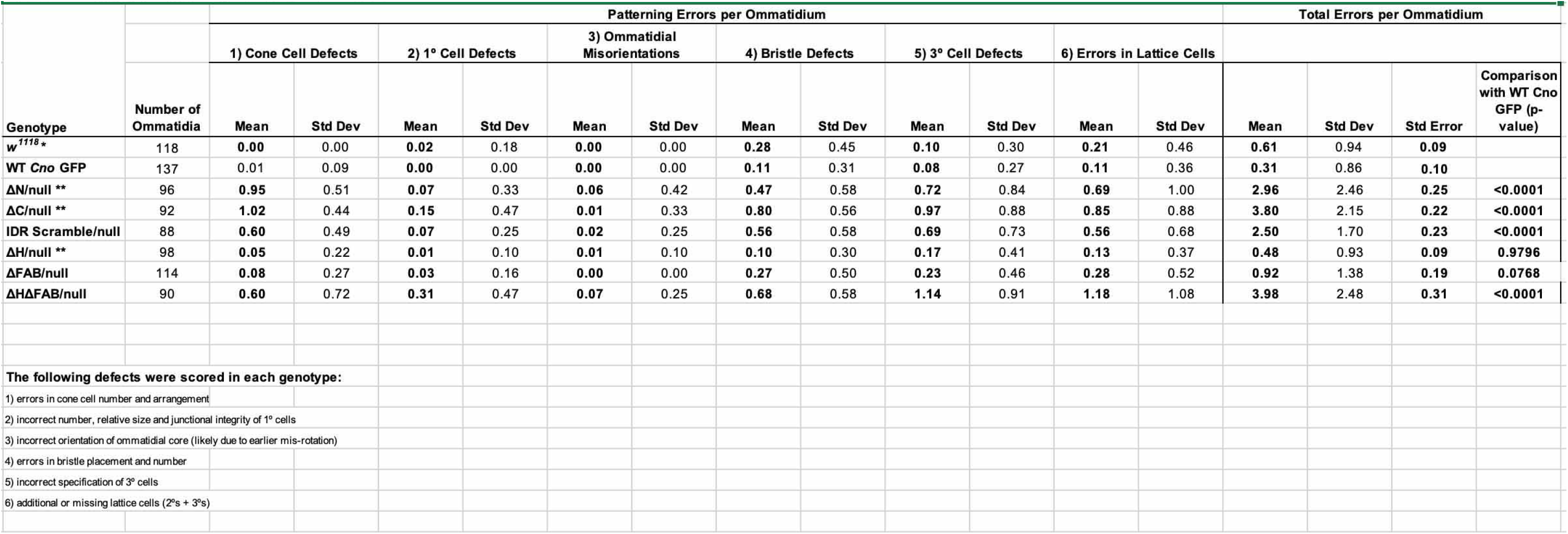
Analyses of Patterning Defects per Ommatidium.

## Notes

### Competing Interest Statement

The authors have declared no competing interest.

### Summary of Updates

Incorrect author order corrected--not sure how they got so scrambled on the initial submission. No other changes were made

## References

Arai, M., Suetaka, S. and Ooka, K. (2024). Dynamics and interactions of intrinsically disordered proteins. Curr. Opin. Struct. Biol. 84, 102734.

Boettner, B., Harjes, P., Ishimaru, S., Heke, M., Fan, H. Q., Qin, Y., Van Aelst, L. and Gaul, U. (2003). The AF-6 homolog canoe acts as a Rap1 effector during dorsal closure of the Drosophila embryo. Genetics 165, 159–169.

Bonello, T. T., Perez-Vale, K. Z., Sumigray, K. D. and Peifer, M. (2018). Rap1 acts via multiple mechanisms to position Canoe and adherens junctions and mediate apical-basal polarity establishment. Development 145, dev157941.

Cermakova, K. and Hodges, H. C. (2023). Interaction modules that impart specificity to disordered protein. Trends Biochem Sci 48, 477–490.

Cheong, H. S. J., Nona, M., Guerra, S. B. and VanBerkum, M. F. (2020). The first quarter of the C-terminal domain of Abelson regulates the WAVE regulatory complex and Enabled in axon guidance. Neural Dev 15, 7.

Cheong, H. S. J. and VanBerkum, M. F. A. (2017). Long disordered regions of the C-terminal domain of Abelson tyrosine kinase have specific and additive functions in regulation and axon localization. PLoS One 12, e0189338.

Choi, W., Harris, N. J., Sumigray, K. D. and Peifer, M. (2013). Rap1 and Canoe/afadin are essential for establishment of apical-basal polarity in the Drosophila embryo. Mol Biol Cell 24, 945–963.

Chong, S. and Mir, M. (2021). Towards Decoding the Sequence-Based Grammar Governing the Functions of Intrinsically Disordered Protein Regions. J Mol Biol 433, 166724.

Chou, T.-B., Noll, E. and Perrimon, N. (1993). Autosomal P[ovo^D1^] dominant female-sterile insertions in *Drosophila* and their use in generating female germ-line chimeras. Development 119, 1359–1369.

Cox, R. T., Kirkpatrick, C. and Peifer, M. (1996). Armadillo is required for adherens junction assembly, cell polarity, and morphogenesis during *Drosophila* embryogenesis. Journal of Cell Biology 134, 133–148.

Devi, B., Nag, N., Uversky, V. N. and Tripathi, T. (2025). Conditional disorder in proteins: functional transitions between order and disorder. Chem Commun (Camb) 61, 16512–16528.

Duan, D., Lyu, W., Chai, P., Ma, S., Wu, K., Wu, C., Xiong, Y., Sestan, N., Zhang, K. and Koleske, A. J. (2023). Abl2 repairs microtubules and phase separates with tubulin to promote microtubule nucleation. Curr Biol 33, 4582–4598 e4510.

Ginell, G. M. and Holehouse, A. S. (2023). An Introduction to the Stickers-and-Spacers Framework as Applied to Biomolecular Condensates. Methods Mol Biol 2563, 95–116.

Gong, R., Reynolds, M. J., Sun, X. and Alushin, G. M. (2025). Afadin mediates cadherin-catenin complex clustering on F-actin linked to cooperative binding and filament curvature. Sci Adv 11, eadu0989.

Gurley, N. J., Szymanski, R. A., Dowen, R. H., Butcher, T. A., Ishiyama, N. and Peifer, M. (2023). Exploring the evolution and function of Canoe’s intrinsically disordered region in linking cell-cell junctions to the cytoskeleton during embryonic morphogenesis. PLoS One 18, e0289224.

Harris, T. J. and Peifer, M. (2004). Adherens junction-dependent and -independent steps in the establishment of epithelial cell polarity in Drosophila. J Cell Biol 167, 135–147.

Holehouse, A. S. and Alberti, S. (2025). Molecular determinants of condensate composition. Mol Cell 85, 290–308.

Holehouse, A. S. and Kragelund, B. B. (2024). The molecular basis for cellular function of intrinsically disordered protein regions. Nat Rev Mol Cell Biol 25, 187–211.

Ikeda, W., Nakanishi, H., Miyoshi, J., Mandai, K., Ishizaki, H., Tanaka, M., Togawa, A., Takahashi, K., Nishioka, H., Yoshida, H., et al. (1999). Afadin: A key molecule essential for structural organization of cell-cell junctions of polarized epithelia during embryogenesis. J Cell Biol 146, 1117–1132.

Jensen, C. C., Gurley, N. J., Mathias, A. J., Wolfsberg, L. R., Xiao, Y., Zhou, Z., Bischoff, M. C., Clark, S. E., Slep, K. C. and Peifer, M. (2025). A key role of Canoe’s intrinsically disordered region in linking cell junctions to the cytoskeleton. J Cell Biol 224.

Johnson, R. I. (2021). Hexagonal patterning of the Drosophila eye. Dev Biol 478, 173–182.

Johnson, R. I. and Cagan, R. L. (2009). A quantitative method to analyze Drosophila pupal eye patterning. PLoS One 4, e7008.

Jumper, J., Evans, R., Pritzel, A., Green, T., Figurnov, M., Ronneberger, O., Tunyasuvunakool, K., Bates, R., Zidek, A., Potapenko, A., et al. (2021). Highly accurate protein structure prediction with AlphaFold. Nature 596, 583–589.

Koyama, T., Iso, N., Norizoe, Y., Sakaue, T. and Yoshimura, S. H. (2024). Charge block-driven liquid-liquid phase separation - mechanism and biological roles. J Cell Sci 137.

Kuno, S., Nakamura, R., Otani, T. and Togashi, H. (2025). Multivalent afadin interaction promotes IDR-mediated condensate formation and junctional separation of epithelial cells. Cell reports 44, 115335.

Kunttas-Tatli, E., Von Kleeck, R. A., Greaves, B. D., Vinson, D., Roberts, D. M. and McCartney, B. M. (2015). The two SAMP repeats and their phosphorylation state in Drosophila Adenomatous polyposis coli-2 play mechanistically distinct roles in negatively regulating Wnt signaling. Mol Biol Cell 26, 4503–4518.

Kunttas-Tatli, E., Zhou, M. N., Zimmerman, S., Molinar, O., Zhouzheng, F., Carter, K., Kapur, M., Cheatle, A., Decal, R. and McCartney, B. M. (2012). Destruction complex function in the Wnt signaling pathway of Drosophila requires multiple interactions between Adenomatous polyposis coli 2 and Armadillo. Genetics 190, 1059–1075.

Larue, L., Ohsugi, M., Hirchenhain, J. and Kemler, R. (1994). E-cadherin null mutant embryos fail to form a trophectoderm epithelium. *Proc.* Nat. Acad. Sci. USA 91, 8263–8267.

Lotthammer, J. M., Hernández-Garcia, J., Griffith, D., Weijers, D., Holehouse, A. S. and Emenecker, R. J. (2026). Metapredict enables accurate disorder prediction across the Tree of Life. BioRxiv 10.1101/2024.11.05.622168.

Manning, L. A., Perez-Vale, K. Z., Schaefer, K. N., Sewell, M. T. and Peifer, M. (2019). The Drosophila Afadin and ZO-1 homologues Canoe and Polychaetoid act in parallel to maintain epithelial integrity when challenged by adherens junction remodeling. Mol Biol Cell 30, 1938–1960.

Matsuo, T., Takahashi, K., Suzuki, E. and Yamamoto, D. (1999). The Canoe protein is necessary in adherens junctions for development of ommatidial architecture in the Drosophila compound eye. Cell Tissue Res 298, 397–404.

McGill, M. A., McKinley, R. F. and Harris, T. J. (2009). Independent cadherin-catenin and Bazooka clusters interact to assemble adherens junctions. J Cell Biol 185, 787–796.

McParland, E. D., Butcher, T. A., Gurley, N. J., Johnson, R. I., Slep, K. C. and Peifer, M. (2024a). The Dilute domain in Canoe is not essential for linking cell junctions to the cytoskeleton but supports morphogenesis robustness. J Cell Sci 137.

McParland, E. D., Gurley, N. J., Wolfsberg, L. R., Butcher, T. A., Bhattarai, A., Jensen, C. C., Johnson, R. I., Slep, K. C. and Peifer, M. (2024b). The dual Ras Association (RA) Domains of Drosophila Canoe have differential roles in linking cell junctions to the cytoskeleton during morphogenesis. Journal of Cell Science in press.

Perez-Vale, K. Z. and Peifer, M. (2020). Orchestrating morphogenesis: building the body plan by cell shape changes and movements. Development 147, dev191049.

Perez-Vale, K. Z., Yow, K. D., Johnson, R. I., Byrnes, A. E., Finegan, T. M., Slep, K. C. and Peifer, M. (2021). Multivalent interactions make adherens junction-cytoskeletal linkage robust during morphogenesis. J Cell Biol 220, e202104087.

Pokutta, S., Drees, F., Yamada, S., Nelson, W. J. and Weis, W. I. (2008). Biochemical and structural analysis of alpha-catenin in cell-cell contacts. Biochem Soc Trans 36, 141–147.

Roberts, D. M., Pronobis, M. I., Poulton, J. S., Waldmann, J. D., Stephenson, E. M., Hanna, S. and Peifer, M. (2011). Deconstructing the beta-catenin destruction complex: mechanistic roles for the tumor suppressor APC in regulating Wnt signaling. Mol Biol Cell 22, 1845–1863.

Rogers, E. M., Allred, S. C. and Peifer, M. (2021). Abelson kinase’s intrinsically disordered region plays essential roles in protein function and protein stability. Cell Commun Signal 19, 27.

Rogers, E. M., Spracklen, A. J., Bilancia, C. G., Sumigray, K. D., Allred, S. C., Nowotarski, S. H., Schaefer, K. N., Ritchie, B. J. and Peifer, M. (2016). Abelson kinase acts as a robust, multifunctional scaffold in regulating embryonic morphogenesis. Mol Biol Cell 27, 2613–2631.

Rouaud, F., Sluysmans, S., Flinois, A., Shah, J., Vasileva, E. and Citi, S. (2020). Scaffolding proteins of vertebrate apical junctions: structure, functions and biophysics. Biochim Biophys Acta Biomembr 1862, 183399.

Rubin, A. J., Dao, T. T., Schueppert, A. V., Choi, S., Groves, J. T., Regev, A. and Shalek, A. K. (2026). Disordered protein LAT encodes relative levels of signaling pathways in T cell activation. Science 392, eads6847.

Sawyer, J. K., Choi, W., Jung, K. C., He, L., Harris, N. J. and Peifer, M. (2011). A contractile actomyosin network linked to adherens junctions by Canoe/afadin helps drive convergent extension. Mol Biol Cell 22, 2491–2508.

Sawyer, J. K., Harris, N. J., Slep, K. C., Gaul, U. and Peifer, M. (2009). The Drosophila afadin homologue Canoe regulates linkage of the actin cytoskeleton to adherens junctions during apical constriction. J Cell Biol 186, 57–73.

Schmidt, A., Finegan, T., Haring, M., Kong, D., Fletcher, A. G., Alam, Z., Grosshans, J., Wolf, F. and Peifer, M. (2023). Polychaetoid/ZO-1 strengthens cell junctions under tension while localizing differently than core adherens junction proteins. Mol Biol Cell 34, ar81.

Shibata, H., Toyama, K., Shioya, H., Ito, M., Hirota, M., Hasegawa, S., Matsumoto, H., Takano, H., Akiyama, T., Toyoshima, K., et al. (1997). Rapid colorectal adenoma formation initiated by conditional targeting of the Apc gene. Science 278, 120–123.

Smits, R., Kielman, M. F., Breukel, C., Zurcher, C., Neufeld, K., Jagmohan-Changur, S., Hofland, N., van Dijk, J., White, R., Edelmann, W., et al. (1999). Apc1638T: a mouse model delineating critical domains of the adenomatous polyposis coli protein involved in tumorigenesis and development. Genes Dev. 13, 1309–1321.

Stamos, J. L. and Weis, W. I. (2013). The beta-catenin destruction complex. Cold Spring Harb Perspect Biol 5, a007898.

Stothard, P. (2000). The sequence manipulation suite: JavaScript programs for analyzing and formatting protein and DNA sequences. BioTechniques 28, 1102, 1104.

Sun, D., LuValle-Burke, I., Pombo-Garcia, K. and Honigmann, A. (2022). Biomolecular condensates in epithelial junctions. Curr Opin Cell Biol 77, 102089.

Tanaka-Okamoto, M., Itoh, Y., Miyoshi, J., Mizoguchi, A., Mizutani, K., Takai, Y. and Inoue, M. (2014). Genetic ablation of afadin causes mislocalization and deformation of Paneth cells in the mouse small intestinal epithelium. PLoS One 9, e110549.

Troyanovsky, S. M. (2023). Adherens junction: the ensemble of specialized cadherin clusters. Trends Cell Biol 33, 374–387.

Walther, R. F., Burki, M., Pinal, N., Rogerson, C. and Pichaud, F. (2018). Rap1, Canoe and Mbt cooperate with Bazooka to promote zonula adherens assembly in the fly photoreceptor. J Cell Sci 131, jcs207779.

Wieschaus, E. and Nüsslein-Volhard, C. (1986). Looking at embryos. In Drosophila, A Practical Approach (ed. D. B. Roberts), pp. 199–228. Oxford, England: IRL Press.

Xu, D., Grishin, N. V. and Chook, Y. M. (2012). NESdb: a database of NES-containing CRM1 cargoes. Mol Biol Cell 23, 3673–3676.

Yamamoto, H., Maruo, T., Majima, T., Ishizaki, H., Tanaka-Okamoto, M., Miyoshi, J., Mandai, K. and Takai, Y. (2013). Genetic deletion of afadin causes hydrocephalus by destruction of adherens junctions in radial glial and ependymal cells in the midbrain. PLoS One 8, e80356.

Yang, Y., Mu, J. Y. W. and Lai, L. Protein IDR linkers regulate overall protein conformation and phase separation. Cell Reports Physical Science 6, 102964

Yang, Z., Zimmerman, S., Brakeman, P. R., Beaudoin, G. M., 3rd, Reichardt, L. F. and Marciano, D. K. (2013). De novo lumen formation and elongation in the developing nephron: a central role for afadin in apical polarity. Development 140, 1774–1784.

Yap, A. S., Duszyc, K. and Viasnoff, V. (2018). Mechanosensing and Mechanotransduction at Cell-Cell Junctions. Cold Spring Harb Perspect Biol 10, a028761.

Yap, A. S., Gomez, G. A. and Parton, R. G. (2015). Adherens Junctions Revisualized: Organizing Cadherins as Nanoassemblies. Dev Cell 35, 12–20.

Yu, H. H. and Zallen, J. A. (2020). Abl and Canoe/Afadin mediate mechanotransduction at tricellular junctions. Science 370, eaba5528.

Zhadanov, A. B., Provance, D. W., Jr., Speer, C. A., Coffin, J. D., Goss, D., Blixt, J. A., Reichert, C. M. and Mercer, J. A. (1999). Absence of the tight junctional protein AF-6 disrupts epithelial cell-cell junctions and cell polarity during mouse development. Curr Biol 9, 880–888.

Zheng, Q. H., Zhang, C., Wang, M. X., Xiang, X., Zhang, S., Wang, Y. and Yu, H. H. (2026). Edge-vertex flow enables rapid adhesion reinforcement under tension. J Cell Biol 225.

